# Tweek-dependent formation of ER-PM contact sites enables astrocyte phagocytic function and remodeling of neurons

**DOI:** 10.1101/2023.11.06.565932

**Authors:** Yunsik Kang, Amanda Jefferson, Amy Sheehan, Rachel De La Torre, Taylor Jay, Lucia Chiao, Alex Hulegaard, Megan Corty, Isabelle Baconguis, Zheng Zhou, Marc R. Freeman

## Abstract

Neuronal remodeling generates an enormous amount of cellular debris, which is cleared from the nervous system by glia. At the larva-to-adult transition, *Drosophila* astrocytes transform into phagocytes and engulf degenerating larval synapses, axonal and dendritic debris. Here we show Tweek, a member of the bridge-like lipid transfer protein family, is upregulated in astrocytes as they ramp up their phagocytic function early in metamorphosis, and is essential for internalization and degradation of neuronal debris. Tweek forms a bridge between the endoplasmic reticulum (ER) and plasma membrane (PM), and loss of Tweek disrupts ER-PM contact formation and membrane lipid distribution. Patient-identified mutations in the human homolog associated with Alkuraya-Kucinskas syndrome resulted in similar defects in neuronal remodeling, indicating these are loss of function mutations. We propose Tweek helps establish and maintain ER-PM contacts during astrocyte phagocytic function and drives bulk lipid transfer to the plasma membrane for continued efficient internalization and degradation of neuronal debris.

## Introduction

The nervous system of complex metazoan is initially overbuilt, containing excess neurons and synaptic connections, but is later refined through activity-dependent mechanisms or in response to other developmental cues. The culling of neurons through apoptotic cell death can occur long after neurons have integrated into circuits and can eliminate a large fraction of any given neuronal population. For instance, 50% of chick ciliary ganglion cells undergo cell death after receiving afferent synaptic input and developing axonal projections ^1,2^, and 40% of mouse GABAergic inhibitory cortical neurons die after migrating from their birth site, acquiring cortical inhibitory neuron morphology, and forming synaptic contacts with surrounding cells ^3^. The refinement of axon projections and synaptic contacts can also be extensive in the developing brain ^4–6^. In the dorsolateral geniculate nucleus (dLGN), target cells innervated by retinal ganglion cells (RGCs) initially receive ∼12 synaptic connections from RGCs, but these are later refined to only 2-3 ^7^. While it remains unclear precisely how much remodeling of axonal projections and synaptic contacts occurs across the nervous system of any organism, it is believed to be widespread and involves the elimination of large amounts of neuronal debris. This process is essential for refining the connectivity of immature neural circuits and optimizing their function in adult animals.

Mammalian microglia ^8^, astrocytes ^9,10^, and more recently, oligodendrocyte precursor cells ^11,12^, have been implicated in actively reshaping immature neural circuits in development through their phagocytic activity. Microglia are the resident immune cells in the brain and engulf cell corpses ^13–15^ along with axonal and synaptic debris ^16^. Astrocytes, which are non-professional phagocytes, perform similar roles ^9,17^, and these two cell types appear to act together to sculpt final connectivity in some circuits ^10,18^. A number of molecules have been identified that are expressed in glia that modulate elimination of neuronal cell corpses or axonal and synaptic debris during development. The complement factor C1q is thought to bind to RGC synapses destined for removal in dLGN, and bind the complement receptor CR3 in microglia ^19,20^. Astrocytes mediate synapse elimination through MEGF10 and MERTK in the dLGN ^10,18^. Fractalkine, released from thalamocortical(TC) neurons and microglia, enables microglial CX3CR1 to promote TC synapse elimination in developing barrel cortex ^21^. Microglial Trem2 is required for developmental synapse elimination ^22^ and regulates the ability of astrocytes to eliminate synapses in cortex and hippocampus ^23^. While these studies provide exciting molecular entry points to understand neuron-glia signaling during developmental neuronal remodeling, their precise mechanisms of action remain unclear, and the signaling pathways that enable any glial cell type to recognize, internalize, and degrade cell corpses or neuronal debris remain poorly defined.

Microglia are professional phagocytes and are equipped with signaling pathways used by macrophages to recognize, phagocytose, and degrade engulfment targets ^24,25^. In contrast, astrocytes are non-professional phagocytes, and the cell biology of their phagocytic functions has not been studied in detail. In professional phagocytes, recognition of engulfment targets by phagocytic receptors leads to the extension of the plasma membrane to enclose the particle within a membrane-bound vesicle known as a phagosome ^26^. Following target internalization, nascent phagosomes mature by fusing with early endosomes to form early phagosomes, then with late endosomes resulting in late phagosomes, and finally with lysosomes to form phagolysosomes ^27^. During this maturation process, the phagosomal lumen becomes a highly acidic and oxidizing environment, rich in hydrolytic enzymes capable of digesting its contents ^28^. Astrocytes in both *Drosophila* and mammals express the Draper/MEGF10 receptor, which is required for clearance of cell corpses, axonal and synaptic debris ^17,18^. In *Drosophila* and mammals, internalized debris eventually appears in acidified compartments, likely for degradation similar to events in macrophages ^17,18,29^. However, we have only a superficial understanding of the cell biology of astrocyte phagocytic function, from target recognition through degradation, especially *in vivo*.

In this study, we explored the molecular and cellular biology of astrocyte phagocytosis of neuronal debris during *Drosophila* metamorphosis, and sought to identify those molecules encoded in the fly genome (i.e. among all transmembrane, secreted, or signal transduction molecules) that are required for astrocyte phagocytic function. We report the results of this screen and show that loss of Tweek, a 565kDa protein, is required for astrocytes to efficiently internalize and degrade neuronal debris. Using AI-based structural analysis, we found Tweek is a member of the family of bridge-like lipid transfer proteins (BLTPs), which are components of endoplasmic reticulum (ER)-plasma membrane (PM) contacts sites, and are thought to drive bulk lipid transfer from ER to the plasma membrane. Loss of Tweek disrupts ER-PM contact site formation, and leads to altered distributions of lipids on astrocyte membranes. Finally, we show a patient-derived mutation in *tweek*, which causes Alkuraya-Kucinskas syndrome, leads to defects in ER-PM contact sites and astrocyte phagocytic function.

## Results

### Identification of genes required for glial clearance of neuronal debris

We sought to identify new molecules required for glial-induced neuronal remodeling, and glial recognition, phagocytosis, and digestion of neuronal debris (i.e. cell corpses, axonal/dendritic and synaptic debris). We focused on the peptidergic ventral Corazonin (vCrz)-expressing neurons. At late third instar larval stages (wL3), there are 8 pairs of bilaterally symmetric vCrz neurons (Figure 1A) in the ventral nerve cord (VNC). During *Drosophila* metamorphosis, vCrz neurons undergo apoptosis, and neurites and synapses degenerate, beginning ∼2 hours after puparium formation (APF). By 6 APF, all neurites undergo fragmentation and a small amount of vCrz cellular debris remains, which is completely cleared by head eversion (HE, ∼12 hrs APF) ^30,31^ (Figure 1A). vCrz neurons are of particular interest since glial phagocytosis of even this single cell type is mechanistically compartmentalized: Draper (MEGF10 in mammals) is required for clearance of cell corpses but is dispensable for clearance of neurite/synaptic debris ^17^.

**Figure 1.**
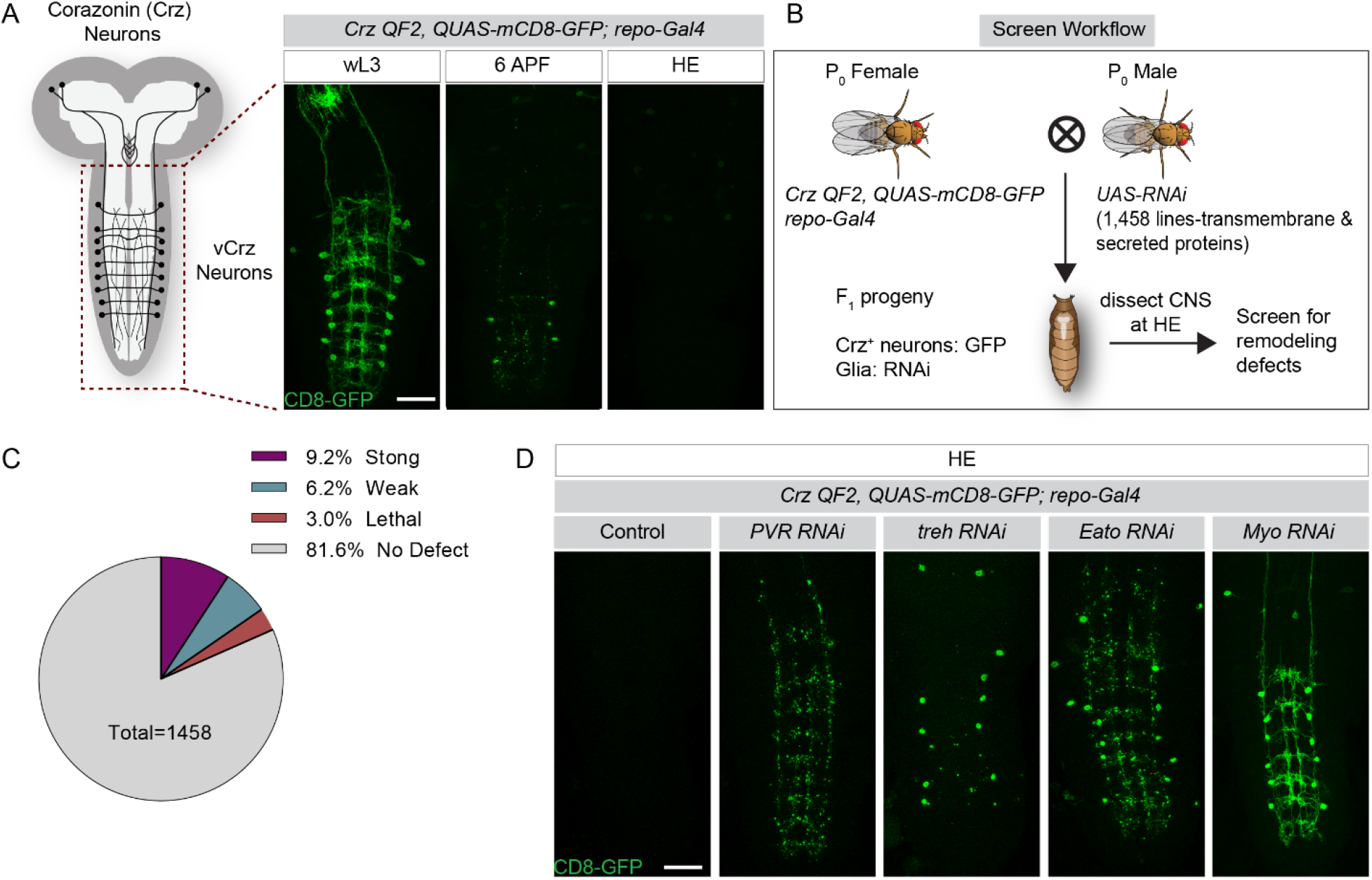
Screen for glial genes required for vCrz neuronal remodeling. (A) Overview of vCrz neuron remodeling - 8 pairs of vCrz neurons in the larval have fragmented and mostly cleared by 6 hours after puparium formation (APF) and are completely cleared at head eversion (HE) (∼12 APF). Unless otherwise stated, all images are maximum Z projections of entire ventral nerve cord (VNC). Scale bar, 50μm. (B) Screen workflow - flies expressing GFP in vCrz neurons (*Crz-QF2, QUAS-mCD8-GFP*) and Gal4 in glia (*repo-Gal4*) were crossed to single *UAS-RNAi* flies, resulting progeny were dissected at head eversion (HE) and assessed for retention of GFP-labeled vCrz debris. (C) Summary of screen results - strong, debris throughout the VNC; weak, some debris at the posterior end of the VNC; lethal, animals that did not survive beyond larval stages. (D) Examples of categories of screen hits - glial knockdown of *PVR* results in only vCrz neurites; *Treh* in only vCrz cell bodies, *Eato* in both vCrz neurites and cell bodies, and *Myo* blocks cell death. Scale bar, 50μm.

**Supplementary Figure 1.**
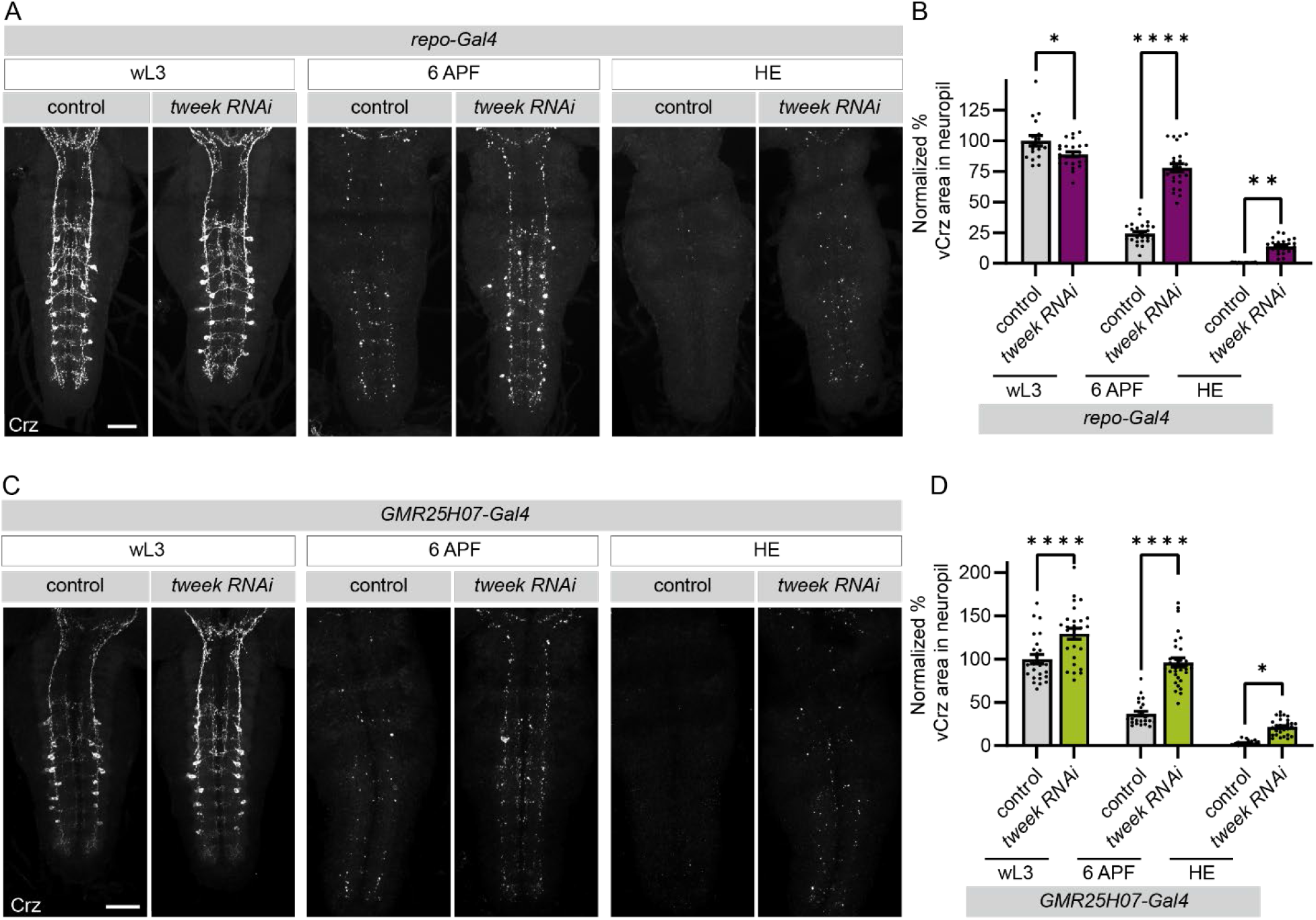
Tweek is required for astrocytic clearance of vCrz neurons. (A) Glial loss of *tweek* suppresses neuronal remodeling. Maximum intensity projection of vCrz neurons labeled with anti-Crz (white). Wandering third instar larvae (wL3); 6 hours after puparium formation (6 APF); and head eversion (HE). Genotypes used were: *repo-Gal4, UAS-nls LacZ* (control), and *repo-Gal4, UAS-tweek RNAiKK101146* (*tweek* RNAi). Scale bar - 50μm. (B) Quantification of vCrz neurite debris area from (A). *repo-Gal4, UAS-nls* (wL3 N=16, 6AFP N=24, and HE N=17)*; and repo-Gal4, UAS-tweek RNAiKK101146* (wL3 N=26, 6AFP N=24, and HE N=24). For all, graphs represent mean ± SEM and each dot represents independent animals. Statistical comparisons were performed using two-way ANOVA with Sidak’s multiple comparisons test. *p<0.05, **p<0.01, ****p<0.0001. Scale bar - 50μm. (C) Astrocyte knockdown of *tweek* suppresses clearance of neurite debris. vCrz neurons labeled with anti-Crz (white). *GMR25H07-Gal4, UAS-nls* (wL3 N=25, 6AFP N=25, and HE N=23), and *GMR25H07-Gal4, UAS-tweek RNAiKK101146* (wL3 N=26, 6AFP N=31, and HE N=31). Scale bar - 50μm. (D) Quantification of vCrz neurite debris area from (C). Genotypes and N as listed in (C). *p<0.05, ****p<0.0001.

To identify glial regulators of neuronal debris clearance, we generated a *Crz-QF2* line to drive membrane-tethered GFP (*QUAS-mCD8-GFP*) in Crz neurons. This allowed us to visualize vCrz neuron remodeling (Figure 1A) while also knocking down genes selectively in glia using a pan-glial driver (*repo-Gal4*) in combination with UAS-driven RNAi lines. We screened a collection of 1,333 genes (1,458 RNAi lines total) representing nearly all transmembrane, secreted, and signaling molecules (i.e. involved in membrane receptor signal transduction pathways) encoded in the *Drosophila* genome (Figure 1B). We assayed for genes that, when knocked down in glia, caused a strong defect in the clearance of vCrz neuronal debris by HE. We identified a total of 125 genes that blocked clearance of vCrz neurites, cell bodies, or both (Figure 1C) (full data set in Supplementary Worksheet 1). These included many known genes, such as *Eato*, an ATPase transporter, whose *C. elegans* homolog *Ced-7* is required for apoptotic corpse clearance ^32^, and mouse homolog *ABCA1*, which is critical for astrocyte phagocytic function after brain injury ^33^ (Figure 1D). We additionally identified Myoglianin, which has been previously shown to encode a glial-released TGFβ molecule that activates the neuronal pruning program^34^. Although many genes required for glial clearance of severed axons were also required for glial remodeling in the pupa (e.g. Draper), to our surprise, several molecules that are known to be required for glial phagocytosis of axonal debris during Wallerian degeneration ^35–42^ did not have a defect in vCrz neuronal debris clearance during metamorphosis (Summary in Supplementary Table 1).

Astrocytes engulf vCrz neurite debris through Draper-independent mechanisms, providing a unique system to identify novel phagocytic molecules or recognition receptors on the astrocyte surface that mediate this event. We, therefore, performed a secondary screen with the astrocyte-specific driver *GMR25H07-Gal4* and assayed 228 high-priority RNAi lines from our initial screen, and those found to cause lethality when driven in all glia with *repo-Gal4*. To control for potential indirect effects on astrocyte phagocytic function due to defects in morphogenesis, we also examined astrocyte morphology at wL3 (by expressing *UAS-mCD8-GFP*). In total, we identified 94 genes that when knocked down by RNAi in astrocytes resulted in significant defects in engulfment of vCrz neuron debris despite having morphologically normal astrocytes (Supplementary Worksheet 1).

### Tweek mutants exhibit defects in clearance of vCrz neurons during neuronal remodeling

We identified *tweek* as a new gene required for astrocyte clearance of vCrz neurons. Tweek protein is very large (565kDa, the 23rd largest in the *Drosophila* genome), and is highly conserved from yeast to humans (29.4% aa identity and 49.6% similarity between fly and human), yet it had no identifiable functional domains by homology searches. Tweek has been proposed to play a role in synaptic vesicle endocytosis by controlling phosphatidylinositol 4,5-bisphosphate {PI(4,5)P_2_} levels at the *Drosophila* NMJ ^43^, and lipid trafficking and cold resistance in *C. elegans* ^44^. Control and *tweek* RNAi animals at larval stages each had 8 pairs of neurons in the VNC, indicating development of vCrz neurons occurred properly in *tweek* RNAi animals. Knocking down *tweek* in all glia with the *repo-Gal4* driver using two independent, non-overlapping RNAi constructs led to strong defects in the phagocytosis of vCrz neuronal debris at HE (Supplementary Worksheet 1). At 6 APF ∼75% of the vCrz cell corpses and neurite debris were cleared in the controls, but in *tweek* RNAi animals only ∼20% had been eliminated (Supplementary Figure 1A, B). By HE all debris was cleared in control animals, while ∼20% of neurite debris was left in *tweek* RNAi animals (Supplementary Figure 1A, B). When we expressed *tweek* RNAi in astrocytes using a strong astrocyte-specific driver (*GMR25H07-Gal4*), we found a similar suppression of clearance of vCrz neurite debris at 6 APF and HE compared to controls (Supplementary Figure 1C, D).

We next used two previously validated null mutants of *tweek* (*tweek^1^* and *tweek*^2^) ^43^ to explore Tweek function. While the majority of the *tweek^1/2^* transheterozygous animals died before metamorphosis, we were able to identify a low number of escapers. In these *tweek^1/2^* animals, larval vCrz neuron morphology was normal (Figure 2A), but *tweek* mutant animals retained ∼50% of the vCrz neurite debris compared to controls at HE (Figure 2B, C). We obtained a BAC clone of *tweek* with a C-terminal mCherry tag that has previously been shown to rescue the lethality of the *tweek* mutant animals ^45^. Introduction of a single copy of this clone completely rescued the vCrz neurite clearance defect (Figure 2B, C), as did resupplying *tweek* specifically in astrocytes with *alrm-Gal4* ^45^(Figure 2B, C). Interestingly, when we assayed for a potential role for *tweek* in other glial phagocytic functions—clearance of olfactory receptor neuron (ORN) axonal debris during Wallerian degeneration—we found no differences in debris clearance between *tweek^RNAi^* and control animals at 1, 3, and 7 days after ORN axotomy (Figure 2D, E).

**Figure 2.**
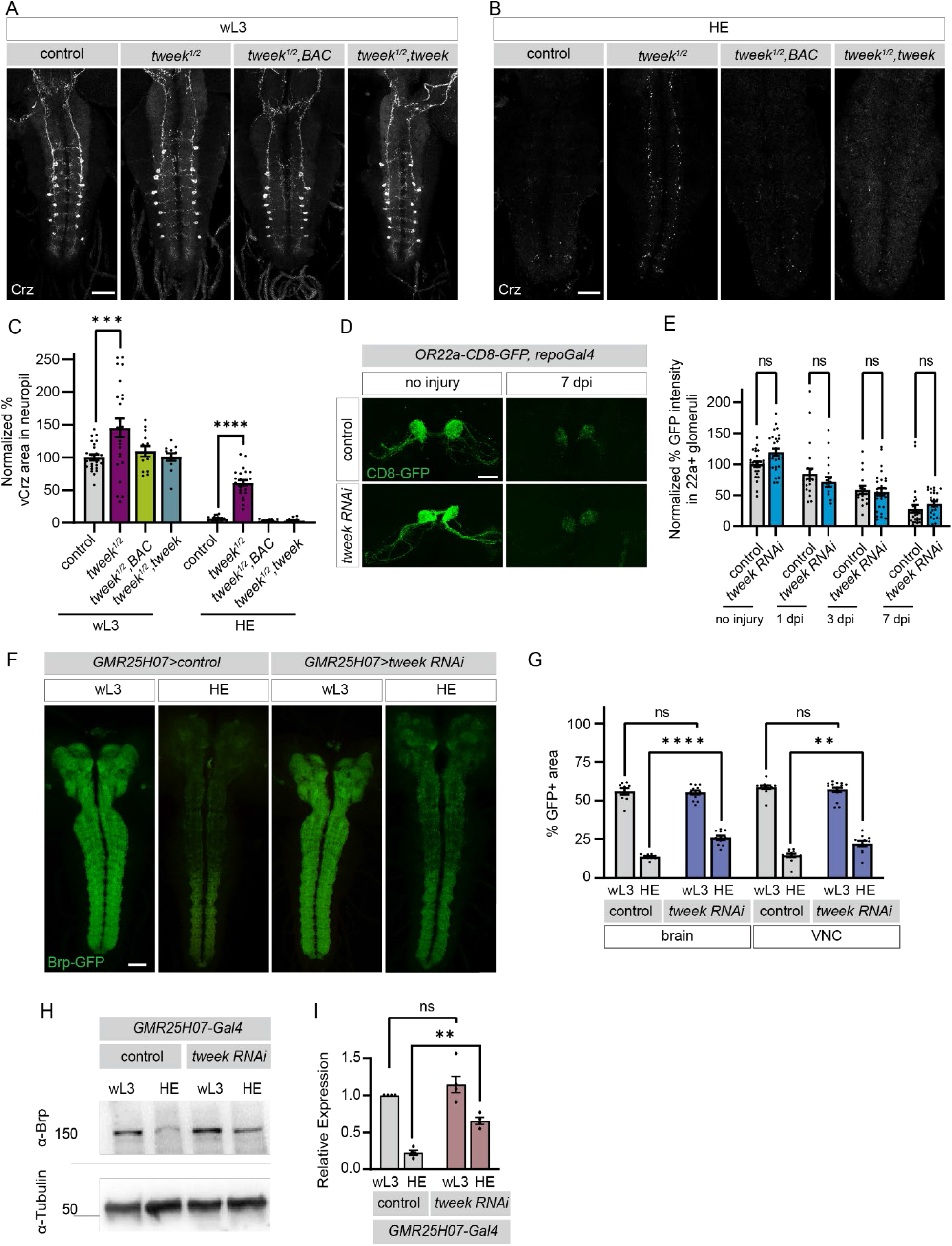
Tweek is required in glia for neuronal remodeling, but not clearance of severed axons. (A) Tweek functions in astrocytes to drive clearance of neurite debris. vCrz neurons labeled with anti-Crz (white) at wL3 and (B) HE. Genotype used were as follows: *w^1118^* (wL3 N=24, HE N=19)*, tweek*^1^*^/^*^2^ (wL3 N=23, HE N=24), *tweek^1/2^; BAC* (wL3 N=13, HE N=19)*, and tweek*^1^*^/^*^2^*; alrm-Gal4, UAS-tweek* (wL3 N=12, HE N=14). Scale bar, 50μm. (C) Quantification of vCrz neurite debris area from (A,B). Statistical comparisons were performed using two-way ANOVA with Sidak’s multiple comparisons test. ***p<0.001, ****p<0.0001. (D) Loss of Tweek does not block glial clearance of severed ORN axons. OR22a^+^ axons labeled with the *mCD8- GFP* in control (*OR22a-GFP, repo-Gal4, UAS-nls-LacZ*) and *tweek RNAi* (*OR22a-GFP, repo-Gal4, UAS-tweek RNAi* KK101146). Glial clearance of axonal debris was assayed with anti-GFP antibody stains (green) after no injury, 1, 3, and 7 days post injury (dpi). Scale bar - 20μm. Genotypes are *OR22a-GFP, repo-Gal4* (no injury: N=13, 1 dpi: N=13, 3 dpi: N=8, 7 dpi: N=14)*, UAS-nls-LacZ* and *OR22a-GFP, repo-Gal4, UAS-tweek RNAi* KK101146 (no injury: N=13, 1 dpi: N=9, 3 dpi: N=14, 7 dpi: N=13). (E) Quantification of axon clearance from the antennal lobe from (D). ns, not significant. (F) Tweek is required in astrocytes for clearance of CNS synapses. Brp-GFP was used to mark presynaptic active zones. Scale bar, 50μm. (G) Brp-GFP+ area within the brain and VNC neuropil were quantified from single z planes. Control-*GMR25H07- Gal4 UAS-nls-lacZ* (N=12 for wL3 and HE), tweek RNAi- *GMR25H07-Gal4 UAS-tweek RNAi* (wL3 N=15 and HE N=12). (H) Western blot analysis of Brp levels in wL3 and HE CNS extracts. Genotypes: *GMR25H07-Gal4, UAS-nls LacZ* (control), G*MR25H07-Gal4, UAS-tweek RNAi* KK101146 (*tweek* RNAi). (I) Quantification of Brp levels in western blots (from F) normalized to Tubulin levels and to control larvae (N=4).

The majority of (and perhaps all) CNS synapses are eliminated during metamorphosis by astrocytes by 48 hr APF ^46^. We examined the role of *tweek* in clearance of synaptic material using a line with an endogenously GFP-tagged *bruchpilot* (*brp*) (Brp-GFP), a marker for presynaptic active zones ^47^. We found Brp-GFP was retained in the VNC and synaptic region of the pupal brain after astrocyte-specific knockdown of *tweek* by confocal microscopy (Figure 2F, G) and Western blots (Figure 2H, I). Together these data support the notion that Tweek is widely used by astrocytes to clear neurite and synaptic debris during neuronal remodeling in the pupa, but is dispensable for glial clearance of axonal and synaptic debris in the context of Wallerian degeneration.

### *tweek* is upregulated in astrocytes as they transform into phagocyte-like cells

We utilized a *tweek* enhancer trap *Gal4* line (*tweek-Gal4*) to examine the spatiotemporal pattern of *tweek* expression during metamorphosis using a nuclear reporter (*UAS-nls-mCherry*). We observed no mCherry+ nuclei in the late larval VNC, suggesting that *tweek* is at most weakly expressed during larval stages (Figure 3A). However, at 2 APF, we observed that all astrocyte nuclei in the VNC were mCherry+ (Figure 3B), and by 6 APF, they were strongly labeled with mCherry (Figure 3 C). While other glial cells expressed *tweek* to some extent, no neuronal expression was observed with this reporter. To further confirm that *tweek* is expressed in astrocytes and to assess its timing of expression, we performed Translating Ribosome Affinity Purification (TRAP) (using *UAS-EGFP-L10a*) with the astrocyte driver *alrm-Gal4* and collected animals at wL3, 2 APF, and 6 APF stages. We immunoprecipitated ribosome-RNA complexes from 150 hand-dissected larval and pupal nervous systems using a high-affinity GFP antibody ^48^, and extracted ribosome-bound RNAs. cDNA libraries were prepared and we performed a quantitative PCR (qPCR) for *tweek. draper* was used as a positive control since it is known to be upregulated in astrocytes during early metamorphosis^17^. We found that *draper* was upregulated ∼7 fold in astrocytes at 2 APF and compared to wL3, as was *tweek* (Figure 3D). Finally, a BAC construct with a mCherry tag at the C-terminal end of Tweek that rescued the phagocytosis defects of *tweek* mutants (Figure 2A-C) was used to examine the expression of endogenous Tweek protein. We observed very little mCherry expression at wL3 stages, but by 6 APF there was significant expression of Tweek in all astrocytes (Figure 3E). Thus, Tweek is highly upregulated in astrocytes at the onset of phagocytic activity.

**Figure 3.**
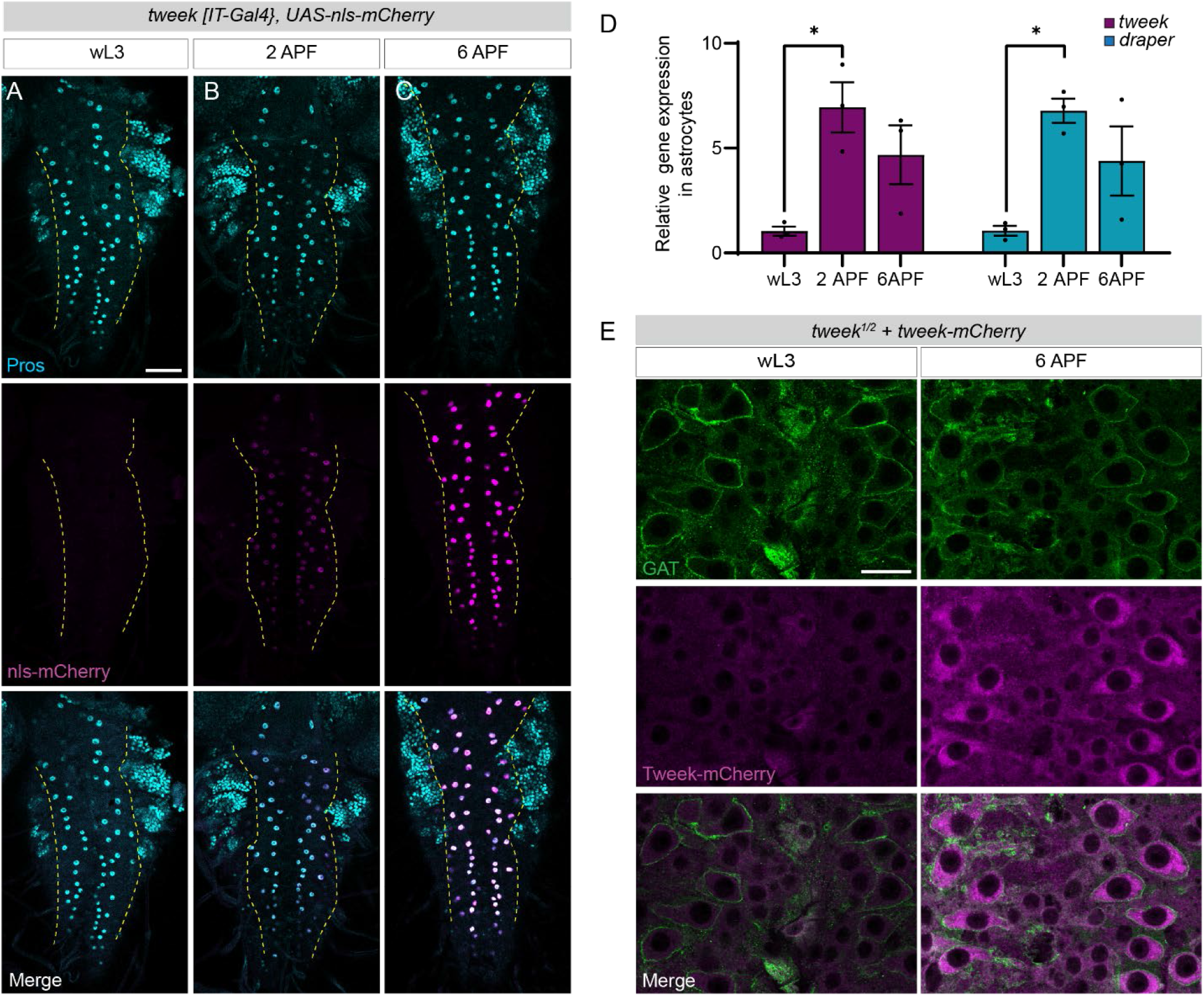
Tweek is expressed in astrocytes as they transform into phagocytes. (A-C) Tweek expression was evaluated in astrocytes within the ventral nerve cord (outlined with yellow dashed line) using an astrocyte nuclear marker (Pros, cyan), and a nuclear mCherry reporter under tweek transcriptional control (magenta, *tweek-Gal4, UAS-nls mCherry*). Scale bar 50μm. (D) TRAP-RNA qPCR graphs of relative expression of *draper* and *tweek* at wL3, 2 APF, and 6 APF. Statistical comparisons were performed using one-way ANOVA with Sidak’s multiple comparisons test. *p<0.05. (F) Endogenously mCherry-tagged Tweek at wL3 and 6 APF. Astrocyte membrane labeled with anti-GAT (green), Tweek-mCherry (magenta). Scale bar, 20μm.

**Supplementary Figure 2.**
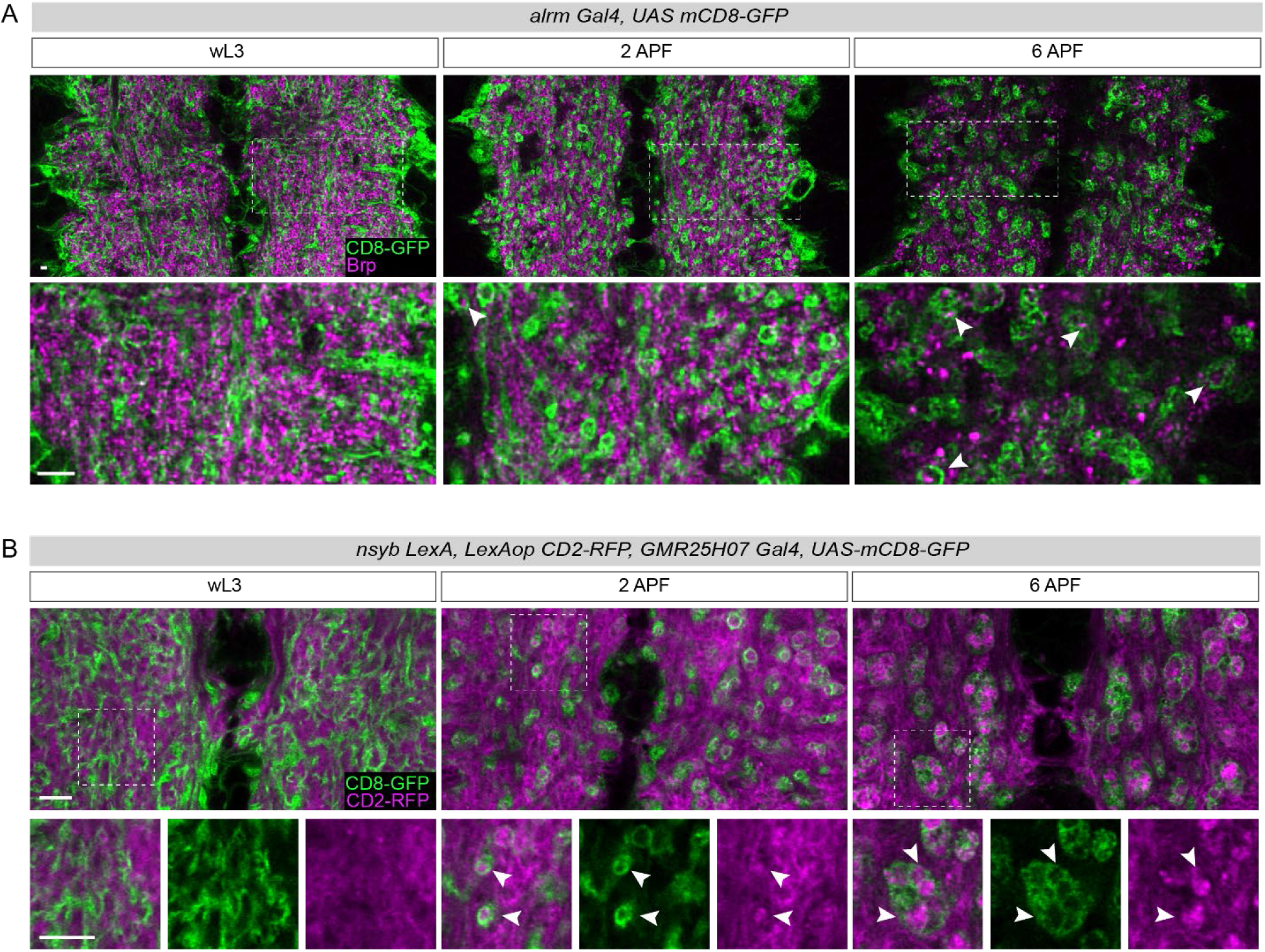
Astrocytes internalize neuronal material. (A) Confocal single Z-plane images of wL3, 2 APF, and 6 APF VNC region in which all astrocyte membrane is labeled with mCD8-GFP and synaptic material stained with anti-Brp (magenta). White arrow heads pointing to internalized synaptic material. Magnification of the boxed region is under each image. Scale bar, 10μm. (B) Confocal single Z-plane images of wL3, 2 APF, and 6 APF VNC region in which all astrocyte membrane labeled with mCD8-GFP and neuronal membrane labeled with mCD2-RFP (magenta). White arrow heads pointing to internalized neuronal membrane. Magnification of the boxed region is under each image. Scale bar, 10μm.

**Supplementary Figure 3.**
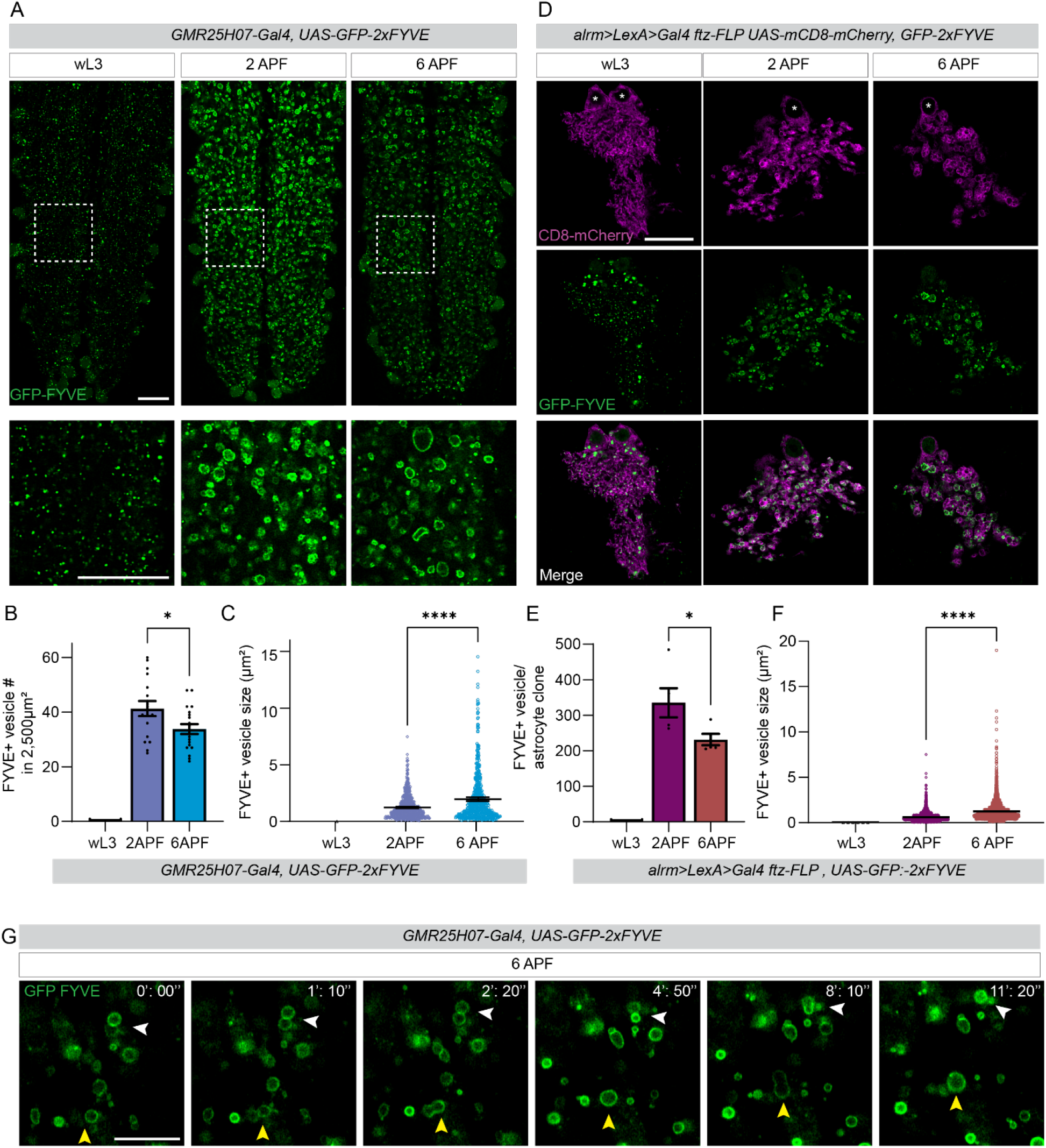
Astrocyte processes form dynamic early phagosomes. (A) Confocal single Z-plane images of wL3, 2APF, and 6APF VNC in which all astrocyte membranes are labeled with GFP-2xFYVE (*GMR25H07-Gal4, UAS-GFP-2xFYVE*). Magnification of the boxed region is under each image. (B) Quantification of GFP-2xFYVE+ vesicles in a 2,500μm^2^ region at wL3, 2 APF, and 6 APF stages. Four different regions of the VNC in 5 independent animals were collected Graphs represent mean ± SEM for (B) and mean ± 95% C.I. for (C) and each dot represents independent imaging region. Statistical comparisons were performed using one-way ANOVA with Sidak’s multiple comparisons test. *p<0.05, ****p<0.0001. Scale bar, 20μm. (C) Quantification of GFP-2xFYVE+ vesicle size in the same region as (B). Each dot represents one vesicle. (D) Single slice confocal single Z-plane images of astrocyte FLP-out clones labeled with mCD8-mCherry (magenta) and GFP-2xFYVE using *alrm>LexA>Gal4 ftz-FLP* in wL3, 2 APF, and 6 APF VNC. (E) Still images of live imaging of GFP-2xFYVE+ vesicles (*GMR25H07>GFP-2xFYVE*). Time is indicated in the top right corner; white and yellow arrowheads point to independent GFP-2xFYVE+ vesicle fusion events.

**Supplementary Figure 4.**
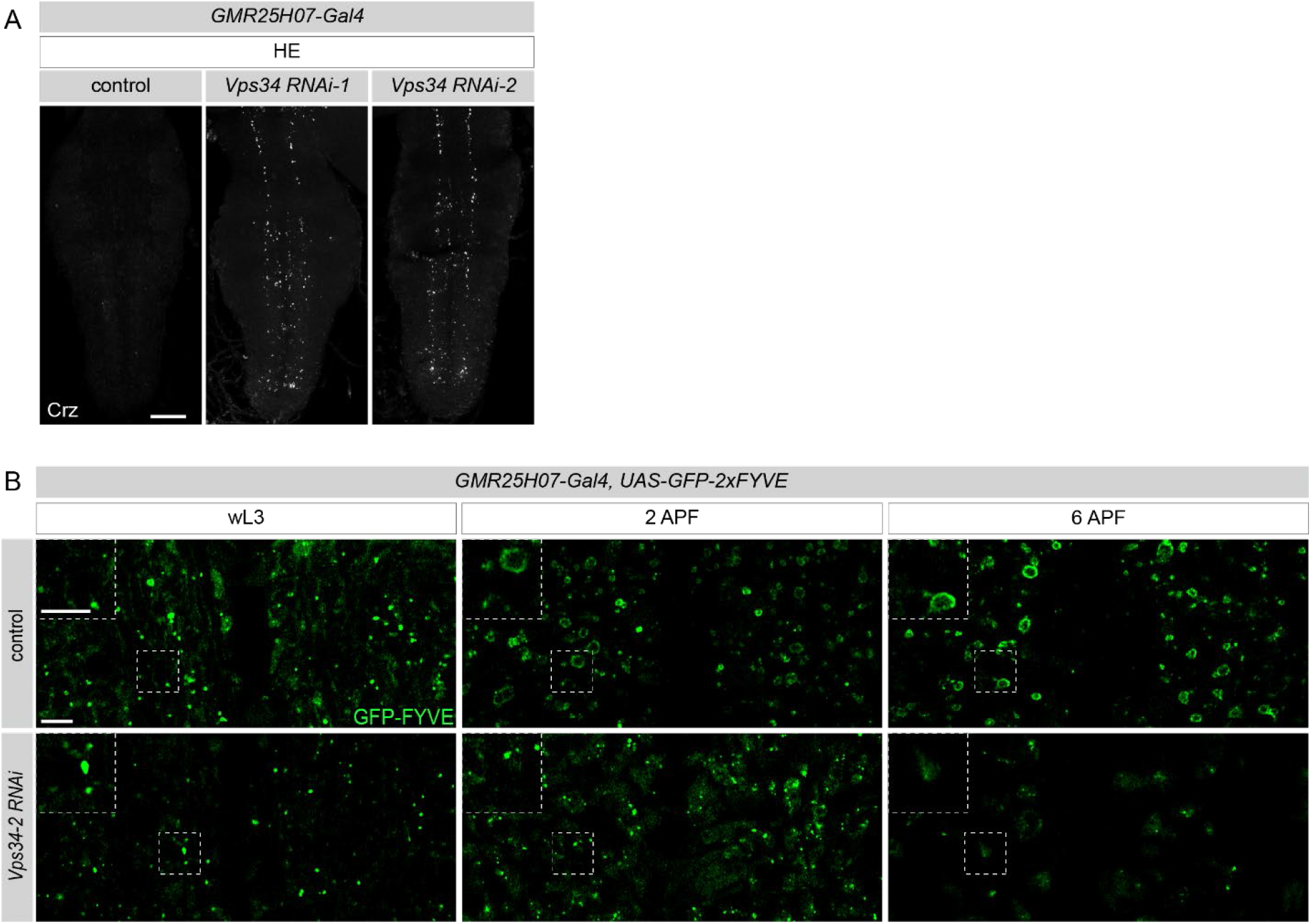
Vps34 is critical for the formation of early phagosomes. (A) vCrz neurons labeled with anti-Crz (white) in pupae at head eversion (HE), maximum z projections. Genotypes: *GMR25H07-Gal4, UAS-nls LacZ, GMR25H07-Gal4, UAS-Vps34 RNAi-1* (TRiP.HMS0026), and *GMR25H07-Gal4, UAS-Vps34 RNAi-2* (TRiP.HMJ30324) Scale bar, 50μm. (B) Confocal single Z-plane images of wL3, 2 APF, and 6 APF VNC in which all astrocyte membrane is labeled with GFP-2xFYVE (*GMR25H07>GFP-2xFYVE*). Magnification of the boxed region is in top left corner of each image. Scale bar, 10μm.

### Astrocytes internalize and digest neuronal debris using canonical phagocytic mechanisms

We previously showed astrocytes transform into phagocyte-like cells at pupariation, where fine processes take on a vesiculated appearance, vesicles become acidified, and a subset housed internalized synaptic material ^17^. To evaluate the time course of engulfment of neuronal material in astrocytes, we labeled neuronal membranes using a membrane-tethered mCherry (*LexAop-mCD8-mCherry*) driven by *nsyb-LexA*, synaptic material with anti-Brp, and astrocyte membranes with *UAS-mCD8-GFP* (driven by *GMR25H07-Gal4*). At 2 APF, synaptic material was found in a small subset of internal astrocyte vesicles, while neuronal membrane (i.e. likely axon/dendrite debris) was found inside nearly all of them (Supplementary Figure 2A, B), and levels of internalized membrane-bound neuronal material and synaptic material increased at 6 APF (Supplementary Figure 2A, B).

To determine whether astrocytes engaged the same phagocytic pathways as professional phagocytes, we first labeled astrocytes using *GFP-2xFYVE* (*GMR25H07>GFP-2xFYVE*), which binds to phosphatidylinositol 3-phosphate (PtdIns3P), a marker for organelles in the endocytic, endocytic recycling pathways, and early phagosomes ^49^. At wL3, GFP-2xFYVE formed small puncta in the astrocyte cell bodies and processes, but by 2 APF GFP-2xFYVE labeled larger, broadly distributed individual vesicles, which increased in number and began to cluster by 6 APF (Supplementary Figure 3A, B). Quantification of GFP-2xFYVE+ vesicles revealed that at 2 APF a larger number of small phagosomes were present in astrocytes, and by 6 APF the total number of FYVE+ phagosomes decreased, but those remaining exhibited an increased volume (Supplementary Figure 3C, D). We speculated that early smaller phagosomes might fuse to generate larger phagosomes. To explore this directly, we modified an *ex vivo* preparation and live imaged astrocytic phagosome dynamics during this developmental window ^50^. To first validate that neuronal remodeling occurred in this preparation, we imaged vCrz neuron cell death and clearance (Movie 1), which occurred properly. When we examined GFP-2xFYVE+ astrocytes, we found GFP-2xFYVE+ vesicles were initially small, but actively fused to increase in size, resulting in larger GFP- 2xFYVE+ vesicles at 6 APF (Supplementary Figure 3E, Movie 2).

To test whether PtdIns3P generation was required for phagocytosis in astrocytes, we knocked down Vps34, which is a class III phosphoinositide 3-kinase that generates PtdIns3P, and inhibition of Vps34 activity leads to defects in phagocytosis by blocking early phagosome formation ^51,52^. To explore its role in astrocytes, we expressed Vps34 RNAi in astrocytes and monitored vCrz neuronal debris clearance and GFP-2xFYVE dynamics. Knockdown of Vps34 led to strong defects in the clearance of vCrz neuronal debris (Supplementary Figure 4A), and in Vps34 knockdown astrocytes, GFP-2xFYVE appeared as puncta at wL3, and by 2 APF or 6 APF, GFP-2xFYVE+ vesicles failed to form (Supplementary Figure 4B), arguing for a blockade of early phagosome formation. We next examined localization of the lysosomal efflux transporter Spinster, which labels lysosomes in *Drosophila* ^53–55^. Unlike GFP-2xFYVE, astrocyte-expressed Spinster-mRFP localized to very few early phagosomes at 2 APF but associated with larger clustered phagosomes at 6 APF, implying that these fused vesicles were likely phagolysosomes (Figure 4A). Indeed, we were able to directly visualize the maturation of GFP-2xFYVE-positive vesicles into Spinster-mRFP-positive vesicles in *ex vivo* imaging assays (Figure 4B, Movie 3). To confirm this, we used the dye LysoSensor to label cellular compartments that have a pH lower than 5, a feature of lysosomes ^56^ and found LysoSensor overlapped with Spinster-mRFP-positive phagosomes, but not with GFP-2xFYVE vesicles (Figure 4A). Together these data support the notion that astrocytes internalize and digest neuronal debris in early phagosomes, these fuse, cluster, and become acidified to digest their internal debris. This stepwise process is quite similar to the cell biology of phagocytosis in professional phagocytes (see Figure 4C for model).

**Figure 4.**
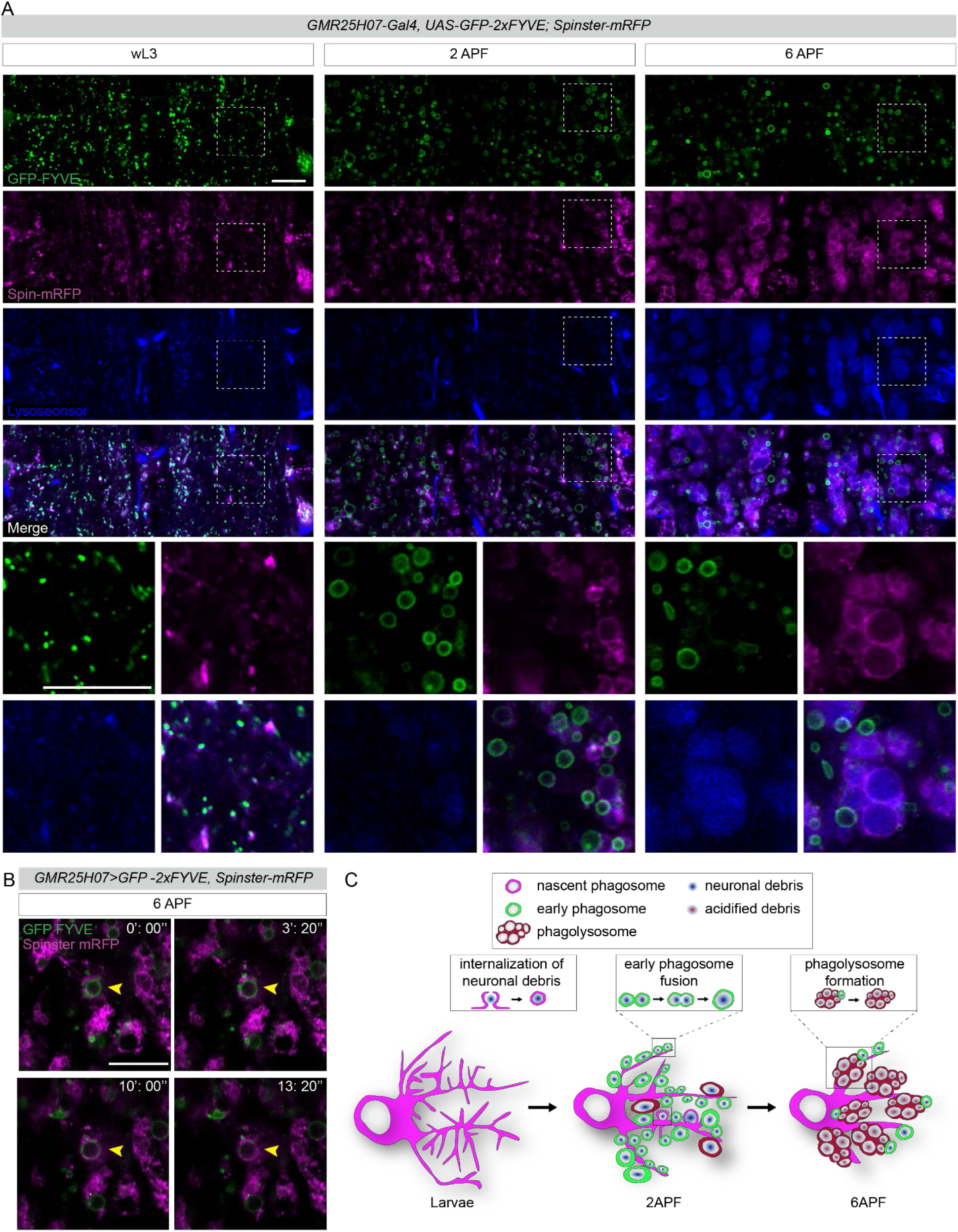
Astrocytes internalize neuronal debris, form early phagosomes that fuse, and ultimately cluster to form phagolysosomes. (A) Confocal single Z-plane images of *GMR25H07-Gal4, UAS-GFP 2xFYVE, UAS-Spinster-mRFP* stained with LysoSensor at wL3, 2 and 6 APF. GFP-2x FYVE in green and Spinster-mRFP in magenta, and LysoSensor in blue. Scale bar, 20μm. (B) Still images of live imaging of GFP-2xFYVE+ and Spinster-mRFP+ vesicles (*GMR25H07-Gal4, UAS-GFP 2xFYVE, UAS-Spinster-mRFP*). Time is indicated in the top right corner, yellow arrowheads point to GFP- 2xFYVE+ vesicle transforming into Spinster-mRFP+ vesicles. (C) Model of how astrocytes execute phagocytosis.

**Supplementary Figure 5.**
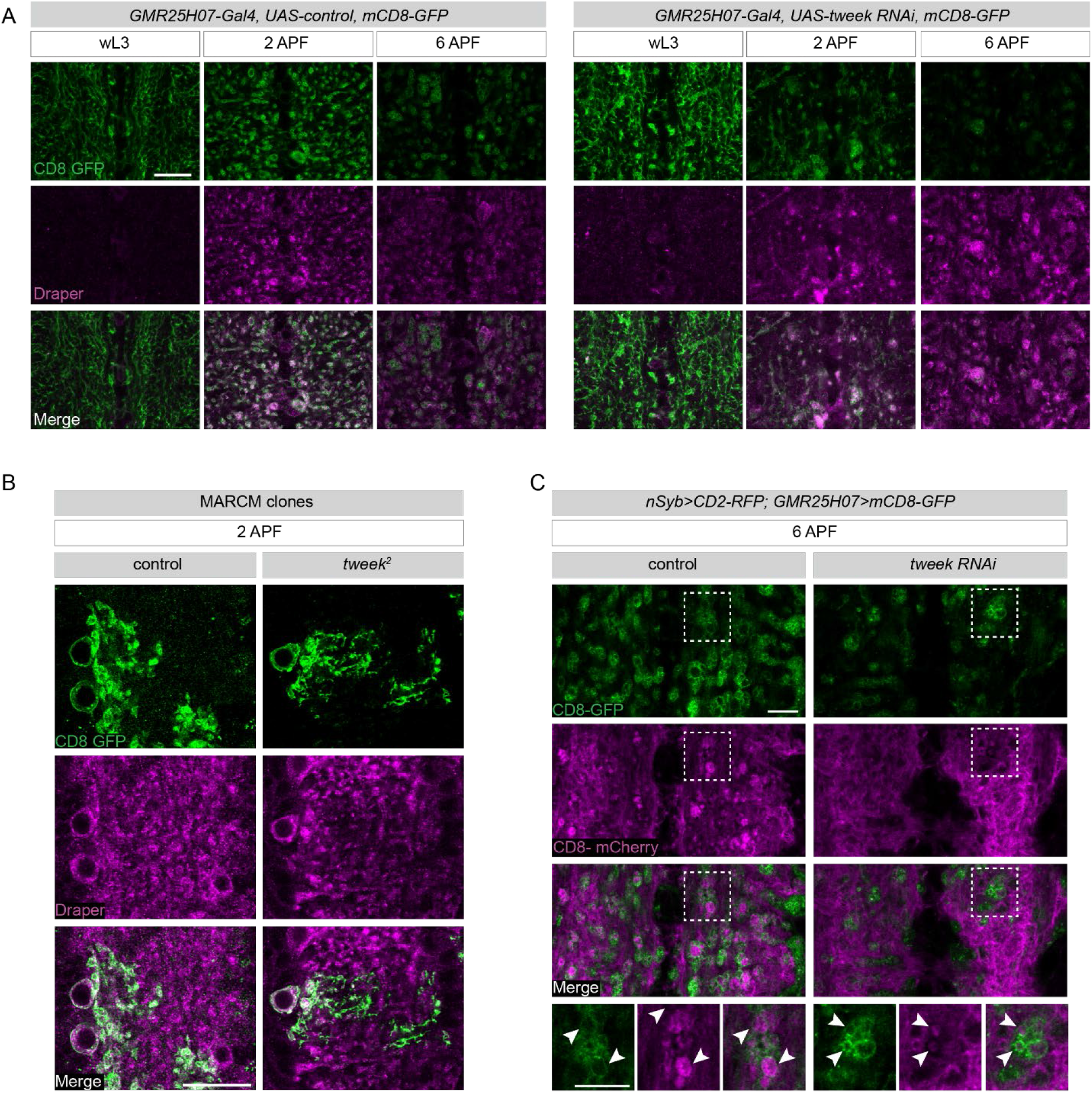
Expression of phagocytic receptor Draper and internalization of neuronal material occurs in *tweek* knockdown astrocytes. (A) Single confocal Z-planes of wL3, 2 APF, and 6 APF in which all astrocyte membrane labeled with mCD8- GFP (*GMR25H07>mCD8-GFP*) stained with Draper (magenta). Scale bar, 20μm. (B) Single Z-plane images of MARCM astrocyte clones of control, tweek mutant (*tweek*^2^) labeled in mCD8- GFP. Phagocytic receptor Draper labeled in Magenta. Scale bar - 20μm. (C) Single confocal Z-plane images of 6 APF VNC region in which all astrocyte membrane labeled with mCD8- GFP and neuronal membrane labeled with mCD2-RFP (magenta). Magnification of the boxed region is under each image. Genotypes are: *GMR25H07>mCD8-GFP, nSyb>mCD2-RFP, UAS-FLP5.DD* and *GMR25H07>mCD8-GFP, nSyb>mCD2-RFP, UAS-tweek RNAiKK101146.* Scale bar, 20μm

### Loss of tweek reduces the capacity of astrocytes to internalize neuronal debris

To explore more deeply how Tweek regulates glial phagocytic function, we first tested whether the loss of *tweek* blocked the early activation of Draper expression. We found in astrocytes expressing *tweek* RNAi and mutant clones, Draper levels were upregulated at 2 APF and 6 APF at levels comparable to controls (Supplementary Figure 5A), suggesting astrocytes have activated their phagocytic program normally. Interestingly, we found that *tweek* mutant astrocytes contained a few small vesicles that contained neuronal debris, but these did not grow in size or number (Supplementary Figure 5B) (Figure 5A, B, C). Surprisingly, at 6 APF early phagosomes appeared to still be forming, although these were significantly reduced in number compared to controls at these time points (Figure 5A, B, C). Moreover, the phagolysosome marker Spinster-mRFP labeled only a small number of clusters of phagolysosomes at 6APF, but they became LysoSensor positive (Figure 5A). These observations are consistent with a model whereby *tweek* mutant astrocytes can initiate the internalization and degradation of a small amount of neuronal debris, but then fail to ramp up this function efficiently. We further confirmed this by examining the ultrastructure of astrocytes at wL3 and 6 APF using transmission electron microscopy. In control astrocytes, several large vesicles filled with cellular debris were present at 6 APF, while in *tweek^RNAi^* astrocytes far fewer such vesicles were present, and they were smaller (Supplementary Figure 6A, B). *tweek* deficient astrocytes can therefore phagocytose and process a small amount of neuronal debris, but appear to have a reduced overall capacity to internalize and process a large amount of neuronal debris generated early in metamorphosis.

**Figure 5.**
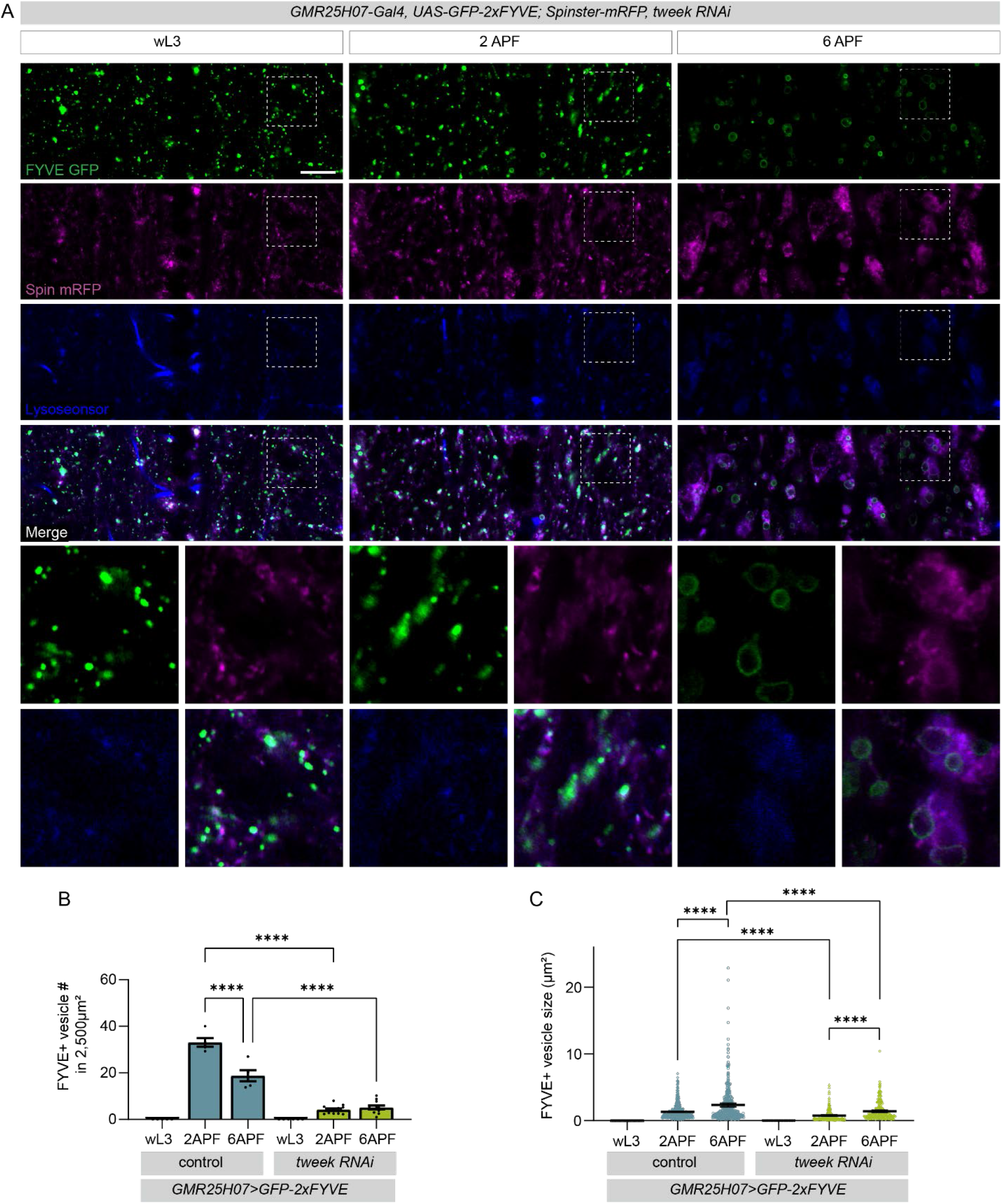
Loss of Tweek decreases the amount of internalized neuronal debris. (A) Confocal Z-plane images of *GMR25H07-Gal4, UAS-GFP 2xFYVE, UAS-Spinster-mRFP, UAS tweek RNAi* stained with LysoSensor wL3, 2 APF and 6 APF. GFP-2xFYVE (green), Spinster-mRFP (magenta), and LysoSensor (blue). Scale bar, 20μm. (B) Quantification of GFP-2xFYVE+ vesicles in a 2500μm^2^ region at wL3, 2APF, and 6APF stages. Four different regions of the VNC were quantified and averaged, represented by a single dot on the graph (control wL3: N=5, control 2 APF: N=5, control 6 APF: N=5. *tweek* RNAi wL3: N=5, *tweek* RNAi 2 APF: N=13, *tweek* RNAi 6 APF: N=11). Genotypes *GMR25H07-Gal4, UAS-GFP 2xFYVE, UAS-Spinster-mRFP, UAS-FLP5.DD* and *GMR25H07-Gal4, UAS-GFP 2xFYVE, UAS-Spinster-mRFP, UAS-tweek RNAiKK101146*. Graphs represent mean ± SEM and each dot represents independent animals. Statistical comparisons were performed using two-way ANOVA with Sidak’s multiple comparisons test. ****p<0.0001. (C) Quantification of GFP-2xFYVE+ vesicle size in the same region as (B). Each dot represents one vesicle.

**Supplementary Figure 6.**
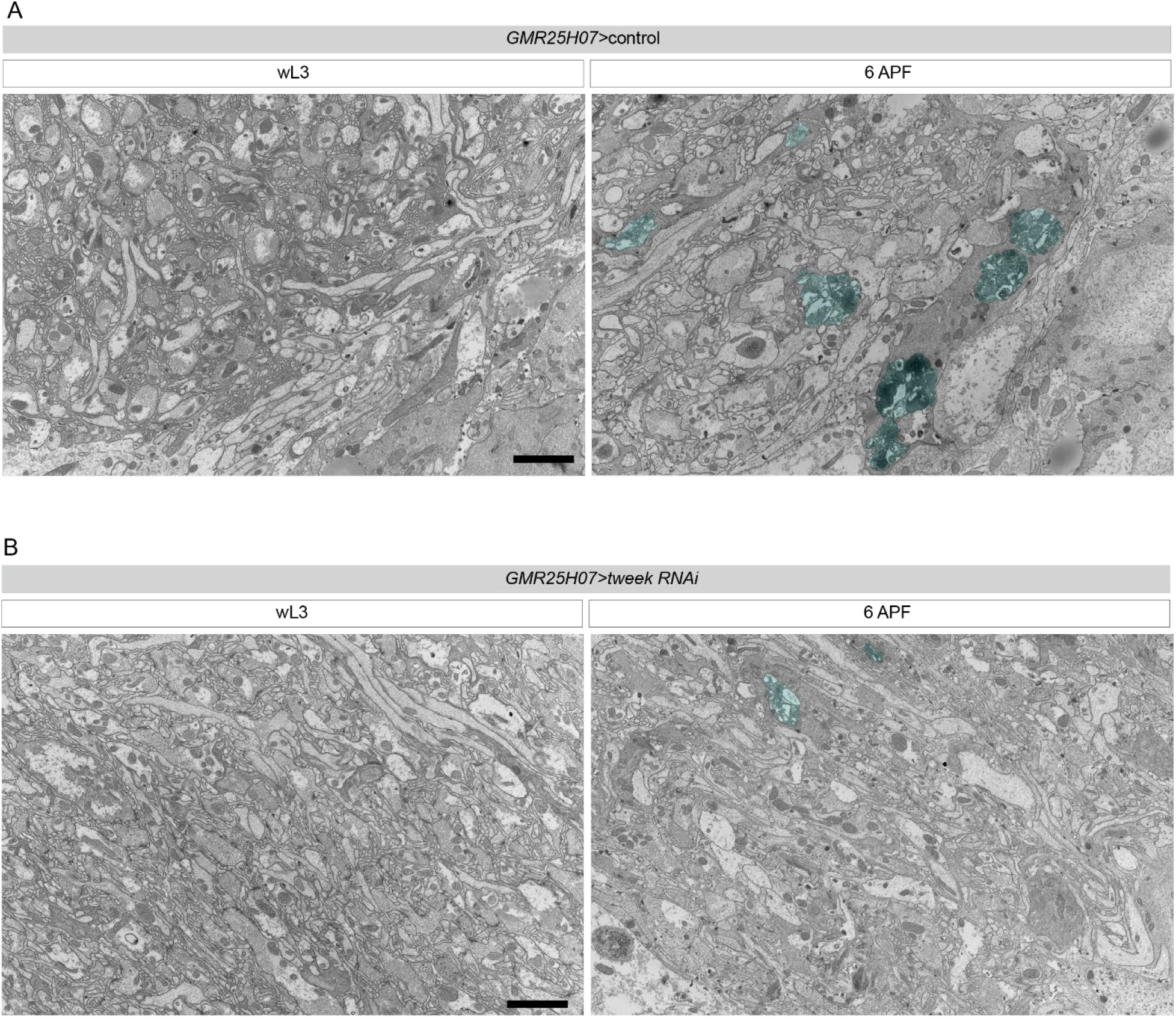
Ultrastructural analysis indicates that *tweek* knockdown astrocytes can internalize small amounts of neuronal debris. TEM image from a cross-section of the abdominal VNC of a (A) control animals and (B) *tweek* knockdown animals (at wL3 and 6APF). Genotype used were:*GMR25H07, UAS-mCD8-GFP* (control)*; GMR25H07, UAS-mCD8-GFP, UAS-tweek RNAiKK101146* (*tweek* RNAi). Cytoplasmic vesicles enwrapped in astrocytic membrane are in turquoise. Scale bar, 2μm.

**Supplementary Figure 7.**
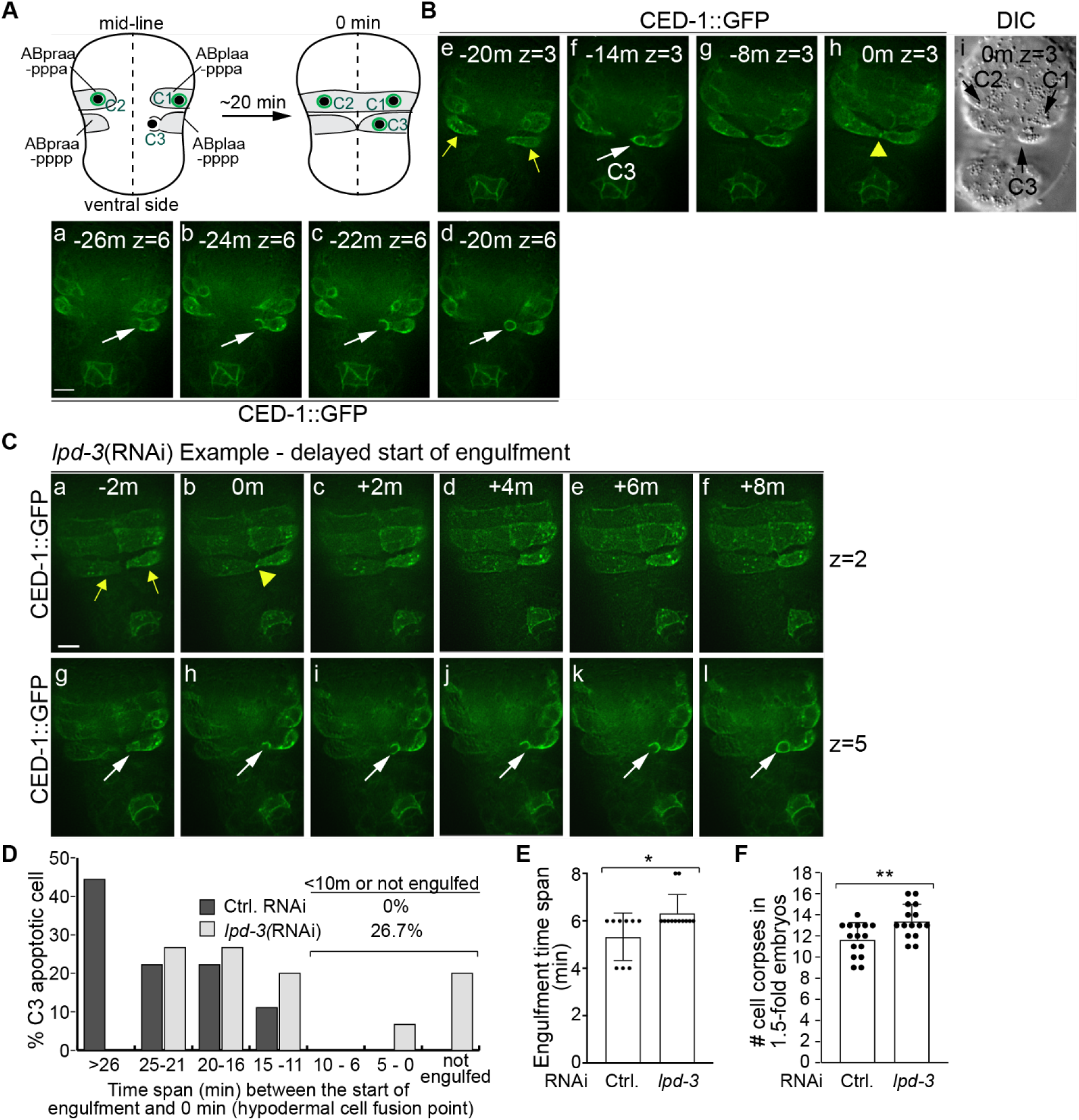
Knockdown of *lpd-3* (tweek C. elegans homolog) does not disrupt enrichment of phagocytic receptors but results in a mild phagocytic defect. (A-C) “0 min” is the first moment when two hypodermal cells ABplaapppp and ABpraapppp come into contact. (A) A diagram illustrating how to monitor the delay of engulfment initiation. Showing here is the ventral surface of a mid-stage embryo. Apoptotic cells C1, C2, and C3 and their designated engulfing cells are marked. “0 min” is the first moment when two hypodermal cells ABplaapppp and ABpraapppp come into contact. In wild-type embryos, the extension of pseudopods from ABplaapppp to C3 occurs on average 22 min before the 0 min time point. (B-C) The engulfment of C3 (white arrows) and the contact point (yellow arrowheads) between ABplaapppp and ABpraapppp (yellow arrows). Time-lapse image series of the ventral side of one each wild-type embryo treated with control RNAi (B) or *lpd-3*(RNAi) (C), respectively. All embryos express P*_ced-1_ ced-1::gfp*, the pseudopod reporter. The “z” numbers indicate the Z-position of the optical section. Scale bars are 5 mm. (D) A histogram of the time spans from the first moment pseudopods were observed to extend towards C3 to “0 min” in wild-type embryos treated with either the empty vector (Ctrl. RNAi) (n=9) or *lpd-3*(RNAi) (n=15). (E) Means and standard deviation (sd) of the time point when engulfment of C3 starts to the moment when C3 is sealed inside a nascent phagosome. n=9 and 12 in “Ctrl. RNAi” and *lpd-3*(RNAi) treated embryos, respectively. Dots are individual numbers. *p<0.05, Student *t*-test. (F) Mean and sd of the numbers of cell corpses counted in 1.5-fold-stage embryos treated with Ctrl. RNAi (n=15) or *lpd-3*(RNAi) (n=15). Dots are values obtained from individual embryos. **p<0.01, Student *t*-test.

**Supplementary Figure 8.**
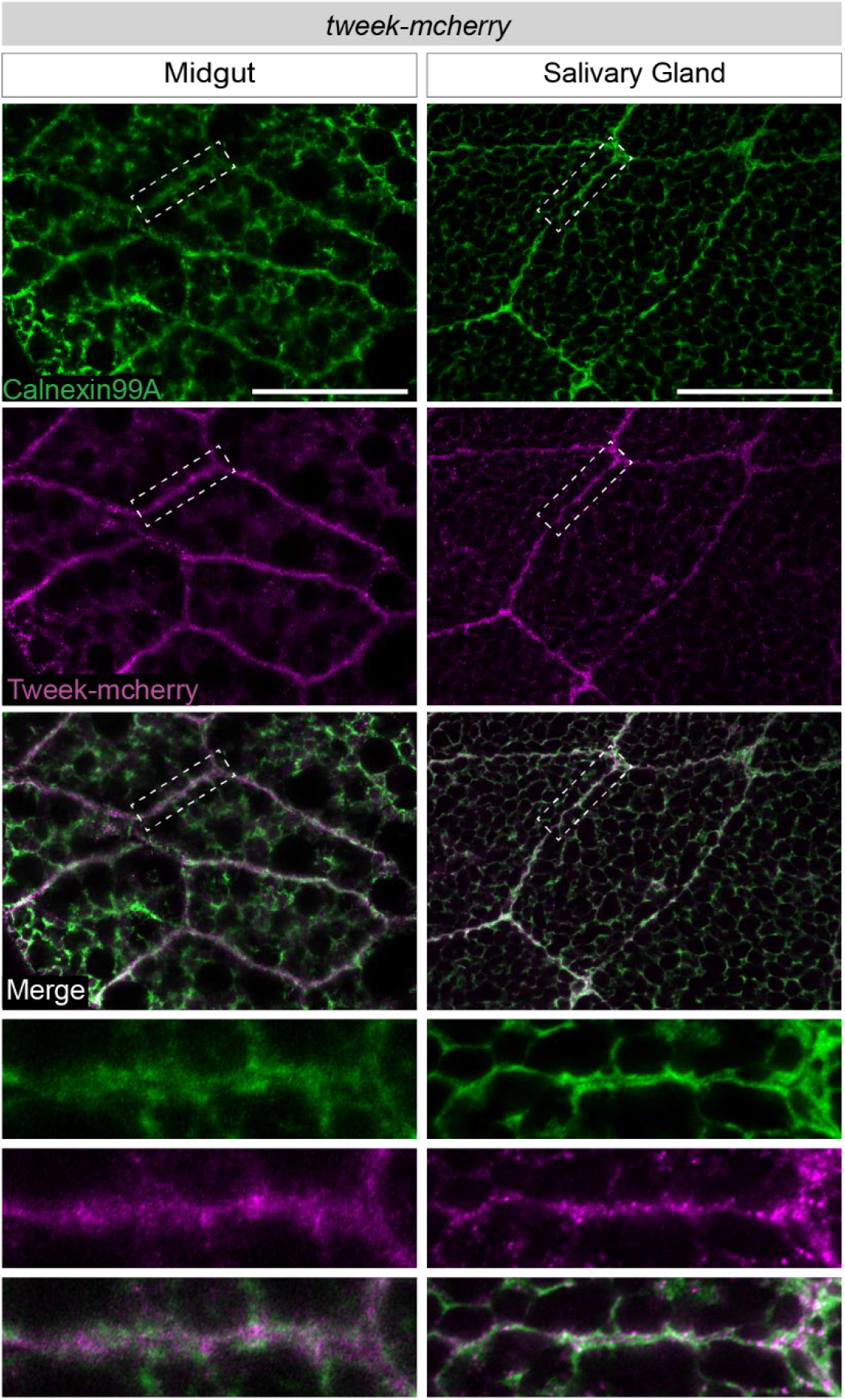
Tweek localizes to the ER and PM in larval salivary glands and midgut. Confocal single Z-plane images of wL3 salivary glands and midgut, stained with ER marker Calnexin99A in green, and endogenous Tweek tagged with mCherry in magenta. Magnification of the boxed region is under each image. Scale bar, 50μm.

**Figure 6.**
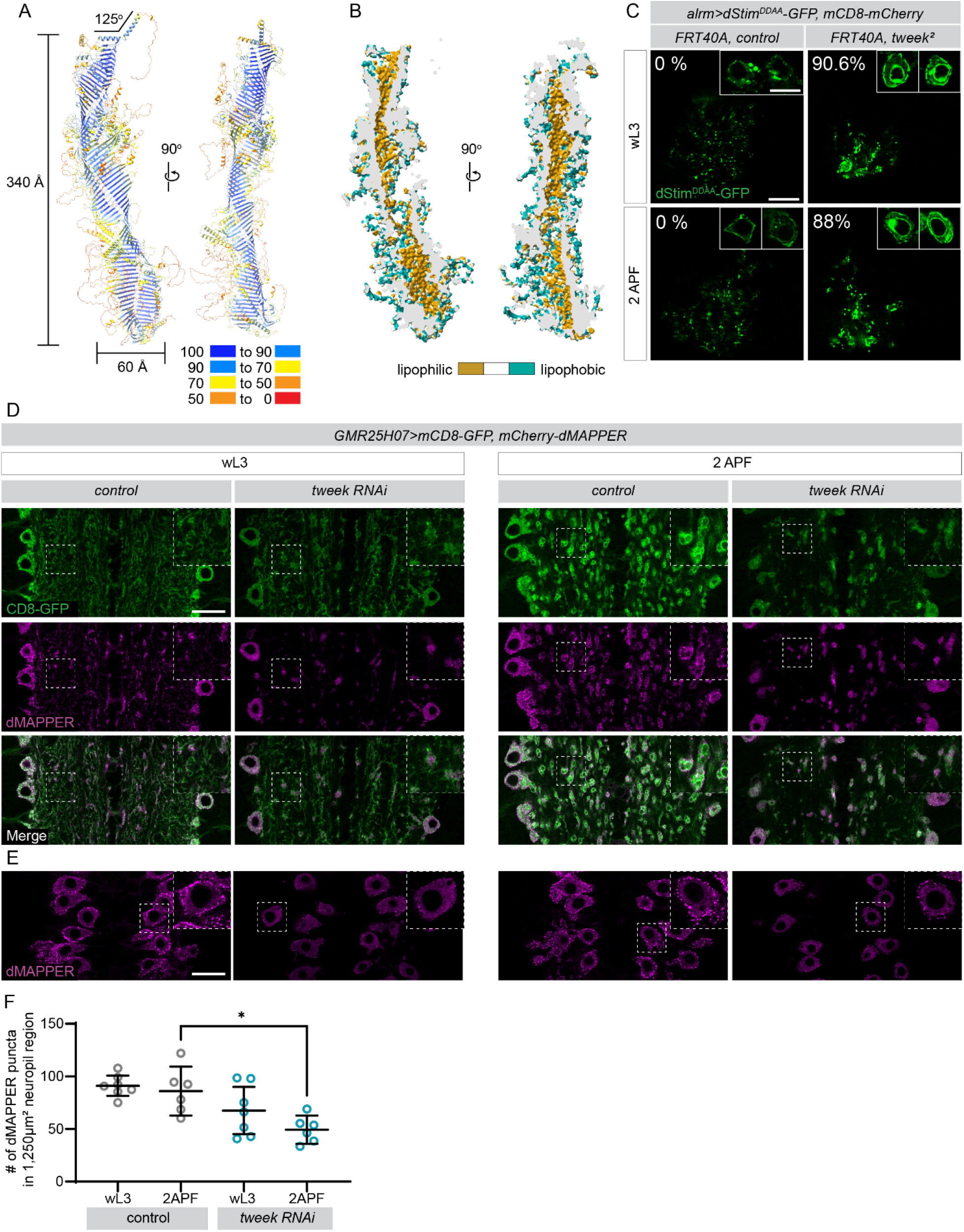
Tweek is required for formation of ER-PM contact sites. (A) Predicted structure of Tweek generated from three structure fragments predicted by AlphaFold v2.0 is shown in cartoon representation. The amino and carboxy termini are located at the top and bottom of the image, respectively. The molecule is colored by model confidence. Regions with scores between 90-100 are expected to have high accuracy, 70-90 with backbone well-modeled, 50-70 with low confidence and therefore caution is advised for model interpretation, and 0-50 should not be considered for analysis. (B) Surface representation of the predicted structure of tweek colored by lipophilicity potential calculated using the program pyMLP as a plugin in ChimeraX. Views of the cross-section of the predicted structure demonstrate a hydrophobic cavity. (C) Single Z-plane confocal images of 2-cell MARCM astrocyte clones of control and *tweek* mutant (*tweek^1^*). Astrocyte membrane is labeled in mCD8-mCherry (magenta) and Stim^DDAA^-GFP (green). Magnification of the astrocyte cell bodies of the clones is on the top right corner of the images. Scale bar - 10μm. Note bright rings of Stim^DDAA^-GFP surrounding nuclei in *tweek* mutants, but not controls. Percentage of Stim^DDAA^-GFP surrounding nuclei are indicated (N>50 nuclei). Scale bar, 10μm. (D) Single confocal Z-planes of wL3 and 2 APF VNCs labeled by mCD8-GFP (green) and the ER-PM contact site marker mCherry-dMAPPER (magenta). Genotypes: *GMR25H07-Gal4, UAS-mCD8-GFP, UAS-FLP5.DD*(control), and *GMR25H07-Gal4, UAS-mCD8-GFP, UAS-mCherry-dMAPPER, UAS-tweek RNAi KK101146* (*tweek* RNAi). Magnification of boxed region in neuropil is on the top right corner. Scale bar, 20μm. (E) View of the cell bodies on the dorsal side the VNC from (D), note ER-PM contact sites (bright magenta puncta) on the cell bodies. (F) Quantification of ER-PM contact puncta in the neuropil. Four different regions of the VNC were collected and averaged to represent a single dot on the graph (control wL3: N=7, control 2APF: N=6, *tweek* RNAi wL3: N=7, *tweek* RNAi 2 APF: N=6). *p<0.05.

### Loss of C. elegans *lpd-3* impairs the clearance of cell corpses but not the enrichment of CED-1

On the plasma membrane of an engulfing cell, the enrichment of phagocytic receptors like Draper/MEGF10 to the site where engulfment is initiated is an important early step in forming phagocytic cups and internalizing engulfment targets. It is possible that Tweek is required for enrichment of Draper to the plasma membrane region in contact with engulfment targets. To determine whether the loss of Tweek altered the enrichment of engulfment receptors to their targets, we examined cell corpse engulfment in *C. elegans*, where one can visualize the enrichment of CED-1 (*C. elegans* Draper/MEGF10) first to the site of engulfing cell-cell corpse contact site and subsequently to the extending pseudopods in live embryos.

We monitored the engulfment of embryonic cell corpse C3 using CED-1::GFP, a GFP-tagged phagocytic receptor that is expressed in engulfing cells and has been observed enriched on the membranes of pseudopods and nascent phagosomes ^57,58^ using an established protocol (Materials & methods). C3 undergoes apoptosis at approximately 320 min post the 1^st^ embryonic cell division and is engulfed immediately (Sup Fig 7A). To determine the moment when the engulfment of C3 starts in a developmental context, we established the moment when ventral hypodermal cells ABplaapppp and ABpraapppp, which extend towards each other during development, make contact, and subsequently fuse with each other at approximately 340 min post the 1^st^ embryonic cell division, as an engulfment-independent reference point (Sup Fig 7A). In wild-type embryos after control RNAi treatment (Materials & methods), the engulfment of C3 was initiated on average 22.7 min before ABplaapppp and ABpraapppp were in contact (Sup Fig 7 B and D). In wild-type embryos after *lpd-3*(RNAi) treatment, we observed stochastic defects in the initiation of the engulfment of C3. Twenty percent of C3 failed to be engulfed by its engulfing cell (Fig D). In some other embryos, the initiation of C3 engulfment is severely delayed (Sup Fig 7 C-D). In addition, there is a mild yet statistically significant longer time period to complete engulfment from an average of 5.3 min to 6.3 min (Sup Fig 7E). Consistent with the defects in engulfing C3, we observed that in 1.5-fold stage embryos (420 min-post the 1^st^ embryonic cell division), *lpd-3*(RNAi) resulted in a modest yet statistically significant increase (14.5%) in the number of cell corpses from 11.7 to 13.4 on average (Sup Fig 7F). This phenotype indicates a decrease in the engulfment efficiency. However, in all embryos treated with *lpd-3*(RNAi), the localization pattern of CED-1::GFP to the plasma membrane of hypodermal cells was normal (Sup Fig 7C(a)). Furthermore, in all cases when C3 was engulfed, regardless of whether the initiation of engulfment was delayed or not, CED-1::GFP is properly enriched on extending pseudopods and subsequently on nascent phagosomes (Sup Fig 7C(g-l)), similar to control embryos (Sup Fig 7A(a-d)). We conclude that LPD- 3 is not required for the proper enrichment of CED-1 to the site of engulfment targets.

**Figure 7.**
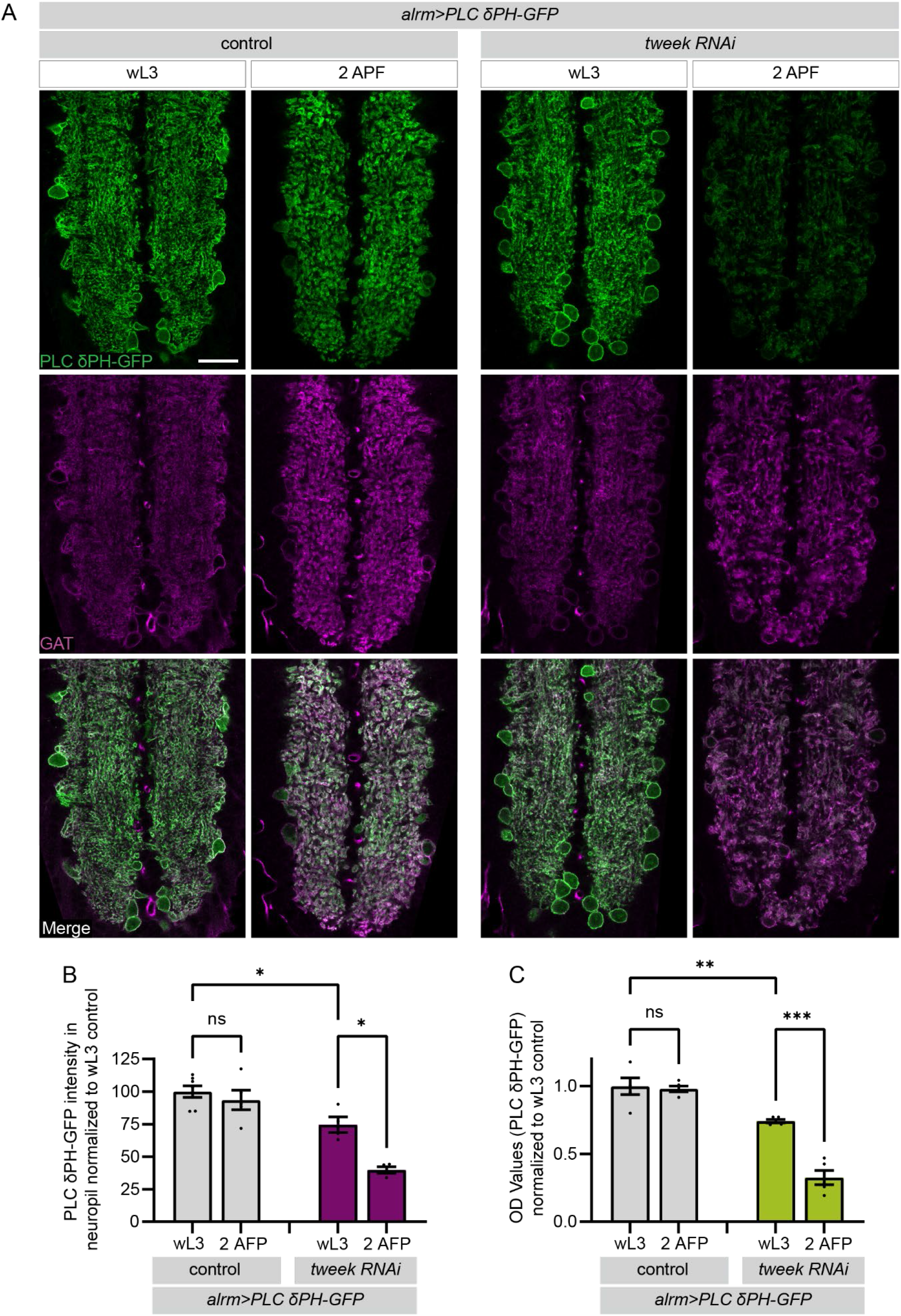
Tweek is required for maintenance of the nascent phagosome marker PtdIns(4,5)P2 during astrocyte engulfment of neuronal debris. (A) Single confocal Z-plane images of wL3, 2 APF, and 6 APF VNC labeled by membrane marker GAT (magenta), and the PtdIns(4,5)P2 biosensor PLCδ-PH-GFP (green). Genotypes: control, *alrm-Gal4,* UAS-PLCδ-PH-GFP*, UAS-FLP5.DD*; and tweek RNAi, UAS-PLCδ-PH-GFP*, UAS-tweek RNAi KK101146.* Scale bar, 20μm. (B) Quantification of PLCδ-PH-GFP intensity in the neuropil. Four different Z-planes of the VNC were collected and averaged to represent a single dot on the graph (control wL3: N=7, control 2APF: N=5, *tweek* RNAi wL3: N=4, *tweek* RNAi 2APF: N=4). *p<0.05. Graphs represent mean ± SEM and each dot represents independent animals. Statistical comparisons were performed using two-way ANOVA with Sidak’s multiple comparisons test. *p<0.05, **p<0.01, ***p<0.001, ns (not significant). (C) Quantification of PLCδ-PH-GFP intensity in the CNS using ELISA (N=5 per genotype).

### Tweek is a bridge-like lipid transfer protein required for formation of ER-PM contact sites during phagocytosis

In an attempt to determine its function, we harnessed the artificial intelligence system AlphaFold2 ^59^ to generate a predicted three-dimensional structure of *Drosophila* Tweek. Because the full-length Tweek is over 5,000 amino acids, we divided the Tweek sequence into three fragments and then assembled a full structure. Our data suggests that Tweek is a remarkably large protein that spans 340 Å at its longest side and 60 Å wide (Figure 6A). Regions of high confidence in prediction primarily reside in the tubular structure consisting of β sheets that assemble in a left-handed twisting fashion. Numerous α helices, ascertained to have well-predicted main chain positions, decorate the β tube and are positioned along β strand interfaces sealing the tubular cavity. The N-terminal segment, comprising an α-helical structure and predicted to span the membrane bilayer, forms a ∼125° bend. Lipophilicity analysis illustrates a hydrophobic core (Figure 6B), in line with the predicted structures of Tweek homologs in yeast (CSF1) and worms (Lpd-3) ^44,60^ (Figure 6B). These structures are highly reminiscent of members of the family of bridge-like transfer proteins (BLTPs), and indeed it has very recently been suggested that Tweek be included as a member of this protein class ^61^. All five members (VPS13, ATG2, Hobbit, Tweek, and SHIP164) localize at membrane contact sites. These proteins are believed to facilitate bulk lipid transfer from one organelle to other organelles via long hydrophobic grooves that are believed to enable vesicle-free lipid transfer ^62^. In line with this possibility that Tweek may serve a similar function, Tweek orthologs in both yeast and worms have been shown to localize to endoplasmic reticulum (ER)-plasma membrane (PM) contact sites ^44,60^.

To assess whether Tweek was similarly localized to these sites in *Drosophila*, we initially assayed its localization in *Drosophila* salivary gland and midgut where the ER and PM are easily visualized due to its large size. We found that Tweek co-localized with the ER marker Calnexin99A, and was also highly enriched at the plasma membrane (Supplementary Figure 8A, B). To monitor ER-PM contact sites in astrocytes, we labeled ER-PM contact sites with Stim^DDAA^-GFP (GMR26H07-Gal4, *UAS-Stim^DDAA^-GFP*) ^63^. We found that Stim^DDAA^-GFP appeared in puncta during wL3 stages in astrocyte processes and around astrocyte cell bodies at 2 APF (Figure 6C). Strikingly, we found in *tweek* mutant MARCM clones, Stim^DDAA^-GFP formed aggregates in the processes of astrocytes, and 90% of all astrocyte cell bodies had Stim^DDAA^-GFP aggregated close to the nucleus (Figure 6C), a feature observed in cells that fail to form ER-PM contacts properly^64^. We next labeled ER-PM contact sites with the ER-PM contact marker mCherry-dMAPPER ^65,66^ and labeled astrocyte membranes with GFP (*GMR25H07- Gal4*, *UAS-mCherry-dMAPPER, UAS-mCD8-GFP, UAS-FLP5.DD*). During larval stages, we found bright puncta that represent ER-PM contact sites localize in the processes of astrocytes and at the cortex of cell bodies (Figure 6D). At 2 APF, astrocyte ER-PM contacts localized to nascent and newly-formed phagosomes and adjacent to the plasma membrane in astrocyte cell bodies (Figure 6D). When we expressed mCherry-dMAPPER in *tweek* knockdown astrocytes (*GMR25H07-Gal4, UAS-mCD8-GFP, mCherry-dMAPPER, tweek RNAi*), ER-PM contacts were significantly reduced in astrocyte cell bodies and in processes in wL3 and 2 APF animals (Figure 6D E). These data indicate that Tweek is necessary for the establishment and maintenance of the ER-PM contact sites and we propose that a primary function of Tweek is to drive bulk lipid transfer from the ER to the plasma membrane to enable astrocyte internalization of neuronal debris.

### Phagosomal membrane phospholipid composition is disrupted in *tweek* mutants

A local increase in phosphatidylinositol 4,5-bisphosphate (PtdIns(4,5)P2} levels is one of the first signaling events that occurs during phagocytosis, with an accumulation of PtdIns(4,5)P2 at the nascent phagosomal membrane^67–69^. We tested whether loss of *tweek* disrupted this process during astrocytic phagocytosis of neuronal debris using the biosensor for PtdIns(4,5)P2 (*UAS-PLCδ-PH-EGFP*) ^43,70^. In control astrocytes at wL3 stages, PtdIns(4,5)P2 levels are detectable and are maintained at 2 APF when early phagosomes are forming (Figure 7A, B). In contrast, we found in *tweek* knockdown astrocytes, PtdIns(4,5)P2 levels were slightly reduced even at wL3, and by 2 APF there was a dramatic decrease compared to control animals (Figure 7A, B). We further confirmed these changes in *tweek^RNAi^* animals using quantitative ELISA assays for GFP using the PLCδ-PH-EGFP reporter at wL3 and 2 APF (Figure 7C) ^71^. These data are consistent with the notion that Tweek is required to supply the appropriate lipids to generate phagosomal membranes from the plasma membrane during astrocyte internalization of neuronal debris.

### Alkuraya-Kucinskas syndrome (ALKKUCS) mutations potently suppress astrocyte phagocytic function

Mutations in the human homolog of *tweek*, *BLTP1* (or *KIAA1109*), were identified as the molecular cause of a rare autosomal recessive disease known as Alkuraya-Kucinskas syndrome (ALKKUCS), which manifests in a combination of physical and neurological deficits ^72,73^. One family of patients were compound heterozygotes, inheriting a late premature stop codon R4375*, and an allele with two missense mutations, N216K and D228G^74^. The asparagine at position 216 is conserved from yeast to humans and corresponds to N239 in *Drosophila tweek* (Figure 8A). In our predicted structure, N239 is located on the solvent-exposed side of the molecule and in the loop that connects two antiparallel β strands, at the juncture of two helices and β sheet (Figure 8B). Such a replacement of a polar asparagine with the positively charged and larger residue lysine would be expected to introduce steric clashes with neighboring residues and change molecular interactions in that region. To determine if this mutation affected Tweek function, we generated a *tweek^N239K^* mutant using genome engineering of the endogenous *tweek* locus, and found this mutation was homozygous lethal and failed to complement the loss of function *tweek* alleles. These mutants exhibited a strong defect in the clearance of vCrz neurons (Figure 8C), exhibited fewer single vesicles in cell clones (Figure 8D), and reduced ER-PM contact sites at 2 APF (Figure 8E). These data indicate that the human N216K mutation is likely a loss of function, and is critical for normal Tweek function *in vivo* (Figure 8C).

**Figure 8.**
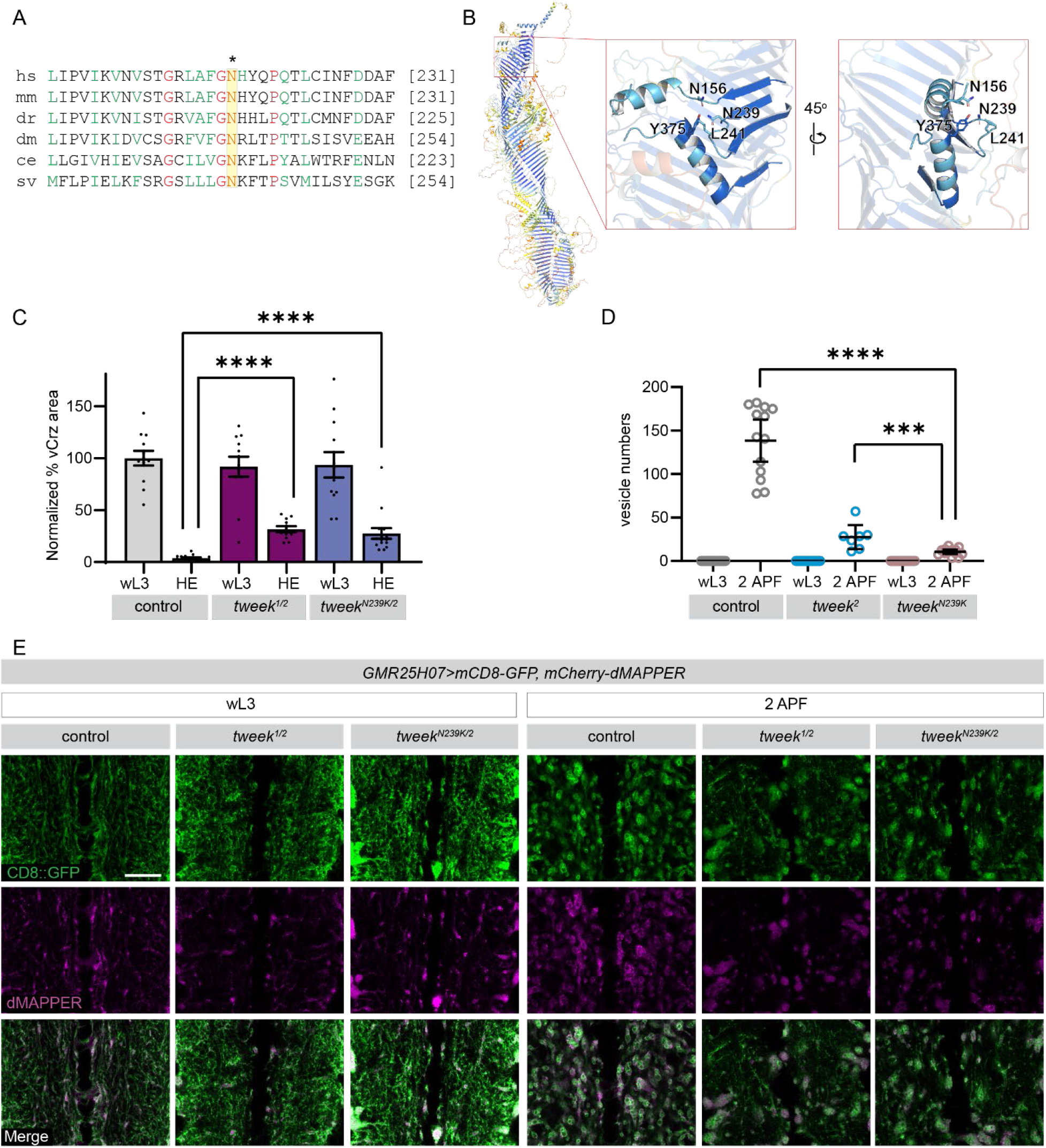
Alkuraya-Kucinskas syndrome (ALKKUCS) mutation potently suppress astrocyte phagocytic function. (A) Amino acid sequence of Tweek homologs in ALKKUCS mutation region. Yellow highlights the conserved asparagine conserved from yeast to humans. (B) The red box near the amino terminal end of the molecule indicates the region in which the residue N239 resides. Close-up view of the red box. The residue N239, shown in sticks representation, resides in a β-hairpin loop and forms multiple interactions between two neighboring helices. Residues from the β-hairpin loop and the two neighboring α helices are shown as sticks to provide molecular context on possible interactions stabilized with N239. (C) Quantification of vCrz neurite debris area from. *w1118* (wL3 N=12 and HE N=17)*; tweek*^1^*^/^*^2^ (wL3 N=12 and HE N=11); and *tweek^N239/2^* (wL3 N=12 and HE N=15). For all, graphs represent mean ± SEM and each dot represents independent animals. Statistical comparisons were performed using two-way ANOVA with Sidak’s multiple comparisons test. ****p<0.0001. (D) Quantification of the number of mCD8-GFP vesicles per astrocyte in MARCM astrocyte clones of control (N=10 for wL3 and N=13 for 2 APF), *tweek* mutant (N=10 for wL3 and N=7 for 2 APF), and ALKKUCS mutation (N=10 for wL3 and N=14 for 2 APF). Graph represents mean ± 95% CI and each dot represents an independent astrocyte. Statistical comparisons were performed using two-way ANOVA with Sidak’s multiple comparisons test. ****p<0.0001. (E) Single confocal Z-planes of wL3 and 2 APF VNCs labeled by mCD8-GFP(green) and the ER-PM contact site marker mCherry-dMAPPER (magenta). Genotypes: control-*GMR25H07-Gal4, UAS-mCD8-GFP, UAS-mCherry-dMAPPER*, tweek mutant-*tweek^1/2^; GMR25H07-Gal4, UAS-mCD8-GFP, UAS-mCherry-dMAPPER,* and disease mutant-*tweek^N239/2^; GMR25H07-Gal4, UAS-mCD8-GFP, UAS-mCherry-dMAPPER.* Scale bar - 20μm.

## Discussion

Astrocytes across species phagocytose neuronal debris, including cell corpses and degenerated axons, dendrites, and synapses ^10,17,18,75^. During pupal neuronal remodeling, *Drosophila* astrocytes engage a stepwise process similar to that found in professional phagocytes: engulfment receptors are recruited to targets, PtdIns(4,5)P2 positive nascent phagosomes form during debris internalization, these mature in a Vps34- dependent manner into PtdIns3P positive early phagosomes that fuse to increase in size, and they mature into late phagosomes that become acidified to phagolysosomes, which then digest debris. While most steps in the phagocytic process appear well conserved, the structure of late phagosomes/phagolysosomes in *Drosophila* astrocytes is unique. Clusters of these structures appear throughout the cell, which to our knowledge have not been described before. Since astrocytes are the primary phagocytes at this developmental time point in the neuropil, and engulf a significant amount of neuronal material, and perhaps also extracellular structures like extracellular matrix, these clusters of phagolysosomes may act as hubs to maximize the efficiency of digestion and recycling of internalized debris.

Our screen sought to comprehensively identify among the majority of transmembrane and secreted molecules, those required in astrocytes for efficient clearance of vCrz neurons. We identified a number of candidates known to play roles in phagocytosis in other contexts (e.g. Draper, Eato, and Ced-7), which validates our screening approach. However, two other themes emerge from our data. First, knockdown or mutation of many genes that are absolutely required for glial engulfment of degenerating axons in other contexts like Wallerian degeneration (e.g. *shark*, *JNK*, *Crk-II*) (Summary in Supplementary Table 1), does not block clearance of vCrz neuronal debris. It is plausible that they are engaged to engulf the debris of other neurons during remodeling that we have not examined yet, but this is striking as the loss of many of these factors can block clearance of axonal debris for weeks after axotomy in the adults ^35–42^. Second, there are no genes that when knocked-out completely block clearance of vCrz neuronal debris (i.e. most show partial clearance by HE). These data argue strongly that the mechanisms used by astrocytes to promote the clearance of neuronal debris during development exhibit a much higher level of genetic redundancy compared to Wallerian degeneration in adults. We note that we only identified one molecule, the TGFβ Myoglianin (Myo), which when knocked down was capable of blocking the initiation of pruning in vCrz neurons, likely by failing to activate the ecdysone receptor (EcR) in vCrz neurons thereby establishing competence to remodel as in other cell types ^76^. Finally, in our screen, genes that had defects in the development or proper morphogenesis of astrocytes resulted in severe phagocytosis defects. This highlights the notion that it is critical to assess the proper development and health of glial cells like astrocytes when evaluating their phagocytic functions.

We identified Tweek as an important modulator of astrocyte phagocytic function. Tweek expression was not detected in larvae but is strongly upregulated as astrocytes transform into phagocytes at pupariation. The small amount of debris that is internalized in *tweek* RNAi animals appears to be processed and digested normally, suggesting the primary role of Tweek may be in facilitating high-capacity phagocytic function. Interestingly, **i**n a screen aimed at identifying regulators of phagocytosis, BLTP1 (human ortholog of Tweek) was found to be critical for efficient phagocytosis of large (but not small) beads ^77^, which also supports the notion it may be engaged during periods of high phagocytic load. Likewise, *C. elegans* Lpd-3 (a homolog of Tweek) was found in a genome-wide screen for suppressors of cell corpse clearance ^78^, and we confirmed that knockdown of Lpd-3 indeed resulted in a mild defect in the clearance of apoptotic cells, seemingly without altering the dynamics of CED-1 (i.e. Draper) localization to cell corpses. Finally, in macrophage cell lines, a CRISPR screen demonstrated that gRNAs targeting BLTP1 led to strong suppression of phagocytosis of the pathogen *Legionella pneumophila*, its primary route of infection ^79^. These studies point to a conserved role for Tweek homologs in phagocytic function. We propose its primary function is to supply the PM with the lipids needed to maintain high levels of internalization of neuronal debris by bulk lipid transfer from the ER.

Tweek is a member of the bridge-like lipid transfer proteins (BLTPs), of which there are five members: Vps13, Atg2, Tweek, Hobbit, and SHIP164 ^61^. While they have low sequence homology, they share a common predicted feature of rotating beta sheets that form a tube-like structure with a long hydrophobic groove ^62^. The BLTPs are thought to promote bulk lipid transfer between membranes ^80^, a process that is essential for the expansion of growing membranes. Where they have been assayed, these proteins localize to endoplasmic reticulum (ER) contact sites, which is a primary source of lipids ^63,80–84^. ATG2 localizes at the ER and autophagosome contact sites in yeast and human cells, where it drives the expansion of autophagosomal membranes, presumably through bulk lipid transfer ^85–88^. Similarly, during *S. cerevisiae* meiosis, VPS13 promotes the expansion of prospore membrane ^89,90^. Deletion of *Atg2* results in a complete blockage of autophagosome formation ^91^, which is reminiscent of our observations that astrocytes fail to generate the normal large population of early phagosomes as they begin to phagocytose neuronal debris during pupal stages.

Precisely how Tweek transfers lipids remains to be determined, but one of the initial membrane phospholipids required for phagosome formation, PtdIns(4,5)P2, is significantly disrupted during early phases of phagocytic function in Tweek mutants. Tweek has also been shown to regulate PtdIns(4,5)P2 at the *Drosophila* NMJ, where its loss perturbs synaptic vesicle endocytosis ^43^, and in worms, Lpd-3 mutants display cold sensitivity, which may be due to their inability to adapt lipid compositions to cold conditions ^92^.

Our work raises an interesting question with respect to the role of Tweek in bulk lipid transfer, during phagocytosis and in other contexts. Our data demonstrates that Tweek is an important structural component of ER-PM contacts in astrocytes, as loss of Tweek disrupts these structures. However, it remains unclear how Tweek is anchored to membranes—although the N-terminus has a transmembrane domain the C-terminus does not. Tweek function in different contexts may be determined by its binding partners. For instance, in yeast sporulation-specific protein Spo71 recruits Vps13 to the prospore membrane ^93,94^, lipid scrambalase Mcp1 to mitochondria ^95^, and the sorting nexin Ypt35 to endosomes, and vacuoles ^96^. Binding partners also likely play critical roles in facilitating BLTP functions in driving membrane formation during lipid transfer. The scramblases Atg9 and XK are essential for the functions of both Atg2 and Vps13, by reorganizing and restoring lipid bilayer symmetry ^97–100^.

Finally, there is a strong association between BLTP mutations and neurological disorders. VPS13A mutations are linked to chorea-acanthocytosis ^101,102^, VPS13B to Cohen syndrome ^103^, VPS13C to Parkinson’s disease with early onset ^104–106^, and VPS13D to spastic ataxia ^107^. Mutations in the human ortholog of Tweek (*BLTP1*) gene have been associated with Alkuraya-Kucinskas syndrome (ALKKUCS), a rare genetic condition characterized by severe intellectual disability, brain abnormalities, arthrogryposis, craniofacial and/or cardiac abnormalities ^72,73,108,109^. In primary dermal fibroblasts from a single patient with ALKKUCS, disruptions in endosomal trafficking and the actin cytoskeleton were observed but the cellular and molecular basis of these phenotypes remained unclear ^74^. We have shown that a single missense mutation found in patients results in a loss of function phenotypes in which internalization of neuronal debris was blocked and ER-PM contact sites were disrupted. This may be attributed to this single mutation causing a complete disruption in protein folding, as it falls in a hinge region, or it may disrupt the binding of essential adaptor proteins necessary for anchoring the BLTP1 to the correct membrane. Future studies of the precise alterations caused by human mutations, and their effect on bulk lipid transfer, should help clarify these issues.

**Table S1.**
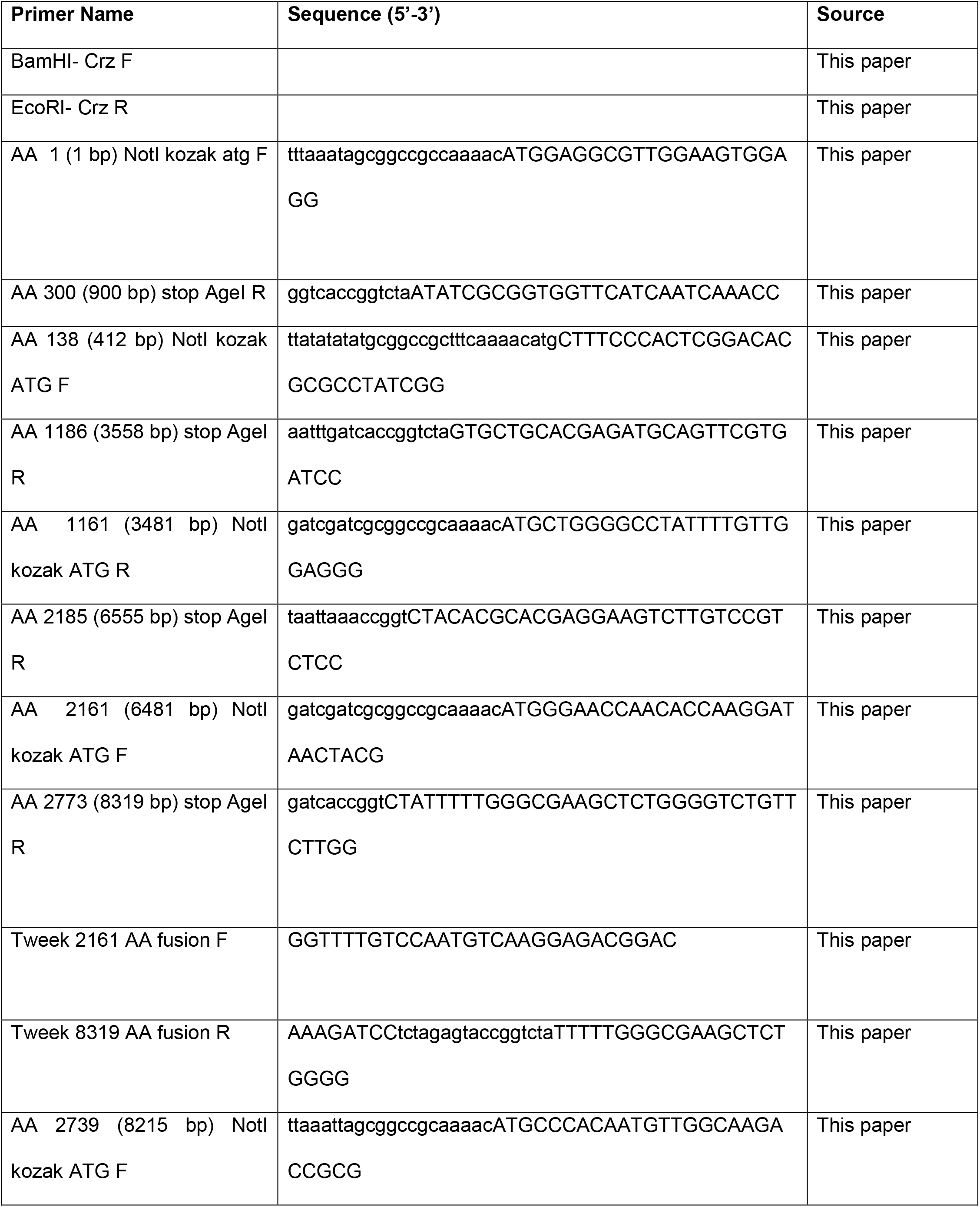
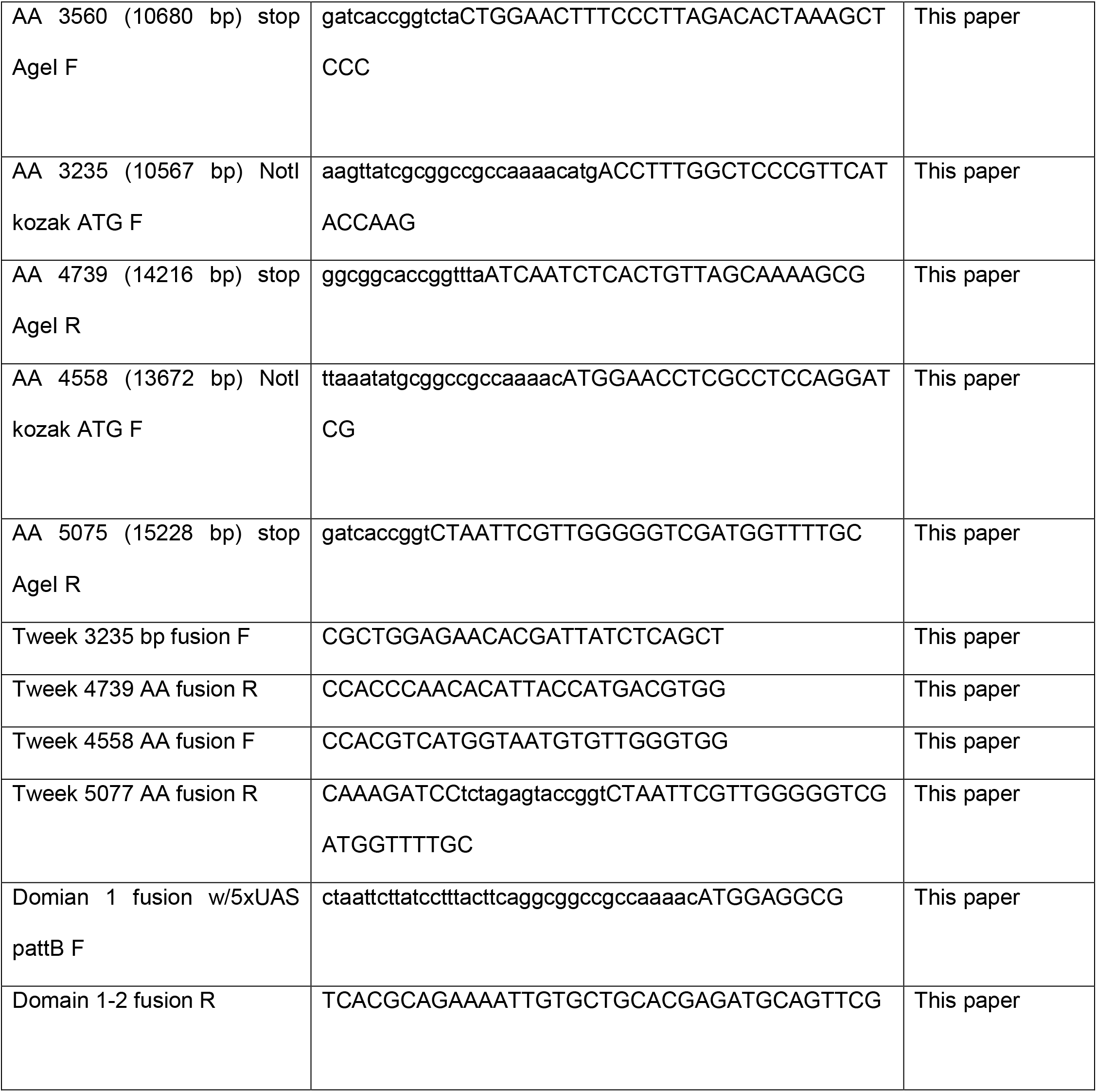
List of primer sequence used in this study, related to Methods section.

## STAR METHODS

## RESOURCE AVAILABILITY

### Lead contact

Further information and requests for resources and reagents should be directed to and will be fulfilled by the lead contacts, Marc R. Freeman.

### Materials availability

Plasmids and transgenic lines generated in this study are available upon request.

### Data and code availability

- All data reported in this paper will be shared by the lead contact upon request.
- This paper does not report original code.
- Any additional information required to reanalyze the data reported in this paper is available from the lead contact upon request.

## EXPERIMENTAL MODEL AND STUDY PARTICIPANT DETAILS

### Drosophila strains

Flies (*Drosophila melanogaster*) raised on standard cornmeal agar with a 12hr/12hr light cycle at 25°C. All experiments were conducted at 25°C. The following lines were used: *w^1118^* (BL3605), *repo-Gal4* (BL7415), *Crz-QF2* (this study), *QUAS-mCD8-GFP* (BL30003), *GMR25H07-Gal4* (Bloomington, 49145), *UAS-tweek^KK101146^ RNAi* (VDRC 110686), *UAS-nls-LacZ* (BL3955), *tweek^1^* (BL39696), *tweek*^2^ (BL39697), *tweek-BAC* ^43^, *UAS-tweek* (this study), *OR22a-GFP* ^110^, *brp^MI02987-GFSTF^* (BL59292), *tweek-IT-Gal4* (BL 63857), *UAS-nls-mCherry* (BL38425), *alrm-Gal4* ^45^, *UAS-mCD8-GFP* ^111^, *nSyb-LexA* (BL52247), *LexAop-CD2-RFP* (BL58755), *UAS-GFP- 2xFYVE* (42712), *UAS-mCD8-mCherry* ^112^, *alrm>nlsLexAfl>Gal4co* ^112^, *UAS-Vps34 RNAi-1* (BL33384), *UAS-Vps34 RNAi-2* (BL64011), *UAS-Spinster-mRFP* (BL42716), *UAS-FLP5.DD* ^113^, *UAS-Stim^DDAA^-GFP* ^63^, *UAS-mCherry-dMAPPER* ^65^, *UAS-PLCδ-PH-GFP* (BL 39693), *tweek^N239K^* (This study).

## METHOD DETAILS

### Generation of *Crz-QF2* stock

To generate the *Crz-QF2* line, we subsequently PCR amplified the genomic DNA sequence spanning the upstream region of the *Crz* gene(–1155 to +78 relative to the transcription start site +1) ^30^ and inserted using BamHI and EcoRI to generate *pattB-Crz-QF2*. The construct was sequence verified by Sanger sequencing (Genewiz) and used for transgenesis (Bestgene), and the resulting stable stock. Primer sequences are described in Table S1.

### Generation of *UAS-tweek* stock

Tweek cDNA is 15,225 bp and was cloned using multiple fragments.

#### pattB-5xUAS Tweek Domain 1 (1-3558 bp)

We generated a cDNA library from RNA collected from w1118 larvae,pupae, and adults using the RNAeasy Plus kit (Qiagen). Using the Superscript III kit First-Strand Synthesis (Invitrogen), we generated a cDNA library. We PCR amplified the cDNA sequence spanning 1-900bp with Not1 and AgeI flanking sites and generated pattB-5xUAS Twk 1-900. Next, we PCR amplified the 412-3558bp region using the cDNA libary flanked by NotI and AgeI and made pattB 5xUS Tweek 412-3558. To make the final pattB- 5xUAS Tweek Domain 1, we cut pattB-5xUAS Tweek 1-900 with NheI and AgeI to remove the end and cut pattB 5xUS Tweek 412-3558with NheI and AgeI gel isolated the 3kb fragment using the Quiagen gel isolation kit and ligated this fragment into Nhe and Age cut pattB-5xUAS Tweek 1-900. All of the ligations were performed using T4 ligase (REF) and transformation was performed in DH5α cells (REF). Primer sequences are described in Table S1.

#### pattB-5xUAS Tweek Domain 2 (3,483- 8,319 bp)

We PCR amplified the 3,483-6,555 bp region from the cDNA library with NotI and AgeI flanking cut sites and generated 5xUAS Tweek 3,483-6,555. We next performed a PCR using the cDNA library to generate a 1,836 bp (6,483- 8,319 bp) fragment with NotI and AgeI flanking and generated 5xUAS Tweek 6,483- 8,319. We performed PCR of the 6,483- 8,319 bp fragment from 5xUAS Tweek 6,483- 8,319. HiFi DNA assmbley was performed (NEB) with 5xUAS Tweek 3,483-6,555 cut with AgeI and a PCR fragment to generate pattB-5xUAS Tweek Domain 2. Transformation was performed in DH5α cells (REF). Primer sequences are described in Table S1.

#### pattB-5xUAS Tweek Domain 3 (8,217- 15,2225 bp)

We PCR amplified region 8,215- 10,680 bp from cDNA library and flanked it with NotI and AgeI to generate 5xUAS Tweek 8,215- 10,680. We repeated the same procedure for 10,567- 14,216 bp. Finally, we repeated the same procedure for region 13,672- 15,225 but used an available cDNA HL01455 (DGRC). We cut 5xUAS Tweek 8,215- 10,680 with AgeI and HiFi DNA assmbley was performed (NEB) using PCR fragments 10,567- 14,216 bp and 13,672- 15,225 bp to generate pattB-5xUAS Tweek Domain. Transformation was performed in DH5α cells (REF). Primer sequences are described in Table S1.

#### pattB-5xUAS Tweek

To generate the final pattB-5xUAS Tweek construct, we cut pattB-5xUAS Tweek Domain 3 with NotI. PCR amplified Domain 1 and PCR Domain 2 and HiFi DNA assmbley was performed (NEB). Transformation was performed in DH5α cells (REF). Primer sequences are described in Table S1.

### Generation of *tweek^N239K^* fly

The *tweek^N239K^* fly was generated following the protocol described in detail ^114^. Briefly, a 4560 bp region containing the DNA sequence for Asparagine at position 239 was selected for swapping to *ywing2+.* Two homology arms were PCR amplified from genomic DNA of *y^1^, w^1118^; attP2{nos-Cas9}/TM6C, Sb, Tb* (Bestgene) flies using Q5 polymerase (NEB), run on a 0.8% agarose gel and purified using the QIAquick Gel Extraction Kit (Qiagen) (Primer sequences in Table S1). The homology arm, pBH donor vector, and *p{y^wing2+^}* were combined by HiFi assembly (NEB). The product was transformed into DH5α Chemically Competent Cells (ThermoFisher) and plated overnight under kanamycin selection. Colonies were cultured for 16 hr at 37°C and DNA was prepared by maxiprep (Qiagen). The entire homology arm sequence and the whole assembled sequence was verified (Genewiz).

sgRNA were cloned into the pCFD3-dU63gRNA plasmid using standard protocols www.flycrisoprdesign.org). Briefly, sense and antisense oligos containing the 20 bp guide target sequence were annealed and phosphorylated with T4 Polynucleotide Kinase (NEB), then inserted between BbsI sites in the plasmid. The ligation product was transformed into DH5α Competent Cells (ThermoFisher) and plated overnight under ampicillin selection. Colonies were cultured, DNA prepared by miniprep (Qiagen), and sequences verified before injection (Genewiz). To generate the *tweek^ywg2+^* flies, we injected a mix of 25 ng/μl sgRNA and 150 ng/μl donor DNA into *y^1^, w^1118^; attP2{nos-Cas9}/TM6C, Sb, Tb* (Bestgene) flies. Once the embryos eclosed, they were crossed to *y,w* flies. The offspring were screened for *yellow+* wings and crossed to the appropriate balancer. We sequence-verified the entire homology arm sequence and adjacent cassettes. Next, to replace the *ywing2+* cassette with the tweekN239K sequence, we first generated the donor DNA with the N239K mutation using HiFi DNA assembly (NEB)(primers in Table S1). The sgRNA was generated as explained above using different 20 bp guide target sequences (Table S1). We injected a mix of 25 ng/μl sgRNA and 150 ng/μl donor DNA into our *y;tweek^wg2+/^CyO; nos-Cas9/TM6B* flies and screened for the loss of the *yellow+* wing. These flies were balanced, and the whole replaced region was sequenced to confirm no additional mutations were generated.

### Generation of Corazonin antibody

Corazonin (Crz)-specific antibody was generated in rabbits by injecting synthetic peptide corresponding to a part of the Crz region (VDPDPENSAHPRLSN) conjugated to Keyhole Limpet Hemocyanin (Genemed synthesis).

### *ex vivo* live imaging

The *ex vivo* live imaging protocol was adapted from a previously published method ^115,116^. To perform live imaging, a series of steps were undertaken to ensure optimal imaging conditions and sample preparation.

#### Media Preparation

A modified Schneider’s *Drosophila* Media was prepared by adding 5 ml of Penicillin-Streptomycin (Gibco) to 500 ml of Schneider’s Drosophila Media (Gibco), followed by filtration. This media was dispensed into 50 ml conical tubes, and up to 45 ml was stored at 4°C for future use.

#### Imaging Setup

Live imaging was conducted using 35×10 mm Petri dishes with a Sylgard base (Dow). To hold samples in place, 24g stainless steel wires (Master wire supply) were bent into three "U" shapes, with two small and one slightly larger "U." The larger wire was inserted into one end of the petri dish, with two long pieces of hair tied to it. To maintain tension on the hair, one of the smaller wire pieces was inserted over the two strands of hair, a short distance from where the hair was tied. On the opposite end of the dish, the second smaller wire was inserted halfway down (ensuring that the wires did not come into contact with the imaging lens) and the hair was wrapped around the wire once. To secure the hair, the remaining ends were tapped to the outside of the dish. It was noted that the hair needed to be re-threaded with new hair after a few uses to ensure proper tension.

#### Day of Imaging

On the day of imaging, a 45 ml tube of the pre-made media stock was removed from 4°C storage. To this media, 5 ml of Fetal Bovine Serum (Fisher) and 125 μl of a 4 mg/ml human insulin stock solution (Gibco) were added. The media was shaken for 20 minutes on a shaker, and then 10 ml of the media was removed and transferred to a new conical tube. The remaining media was stored at 4°C and used within a week. The 10 ml of imaging media was oxygenated for 20 minutes with a very slow flow of oxygen. Once fully oxygenated, 25 μl of a 1 mg/ml 20-hydroxyecdysone solution was addedThe media was shaken for an additional 20 minutes to ensure proper mixing and was ready to use at room temperature.

#### Sample Preparation for Live Imaging

For live imaging, enough imaging media was added to the petri dish to fully cover the sample and wires. The central nervous system of appropriately staged larval or pupal specimens was dissected and carefully placed in the center of the petri dish, between the two strands of hair. The leg, eye, and antennal discs were preserved during dissection. To secure the sample in place, the eye discs were positioned under the top strand of hair, while the leg discs were placed under the bottom strand of hair. Ensuring proper tension on the hair, the sample was held tightly in position. The distance and tension of the hair were adjusted as necessary, and the loose end of the hair hanging off the plate was secured with tape to maintain tension. Depending on the imaging duration, additional media was added to the dish using a dropper to ensure a continuous imaging environment.

### Lysosensor Imaging

Lysosensor dye was employed at a concentration of 1 mM in live imaging media (used for ex vivo live imaging). Animals at the appropriate developmental stage were dissected in live imaging media, transferred to the media containing lysosensor for a 5-minute incubation, and subsequently mounted and imaged immediately.

### Antennae ablation assay

Both antennae of the appropriate adults were surgically removed using forceps. The animals were maintained at a temperature of 25°C both before and after ablation and were dissected on the same day. Whole heads were fixed for 16 minutes in 4% PFA in 0.1% PBST and subsequently washed in 0.1% PBST. The brains were dissected in 0.1% PBST, then fixed for 16 minutes in 4% PFA in 0.1% PBST and washed again with 0.1% PBST. All heads and brains were kept on ice in between dissections. GFP was detected using an antibody at a 1:1000 dilution (Invitrogen).

### Method for structure prediction of tweek

The full-length sequence of Tweek consisted of 5,075 residues. To perform structure prediction using AlphaFold v2.0, the full-length sequence was divided into three ∼2,000-residue fragments with 500 residues overlapping between the fragments ^59,117^. The resulting predicted structures were aligned and merged using Coot to generate the full-length structure ^118^. Loops with low confidence scores and participate in clashes with the neighboring fragments were removed for clarity. Structural images were generated using PyMol and ChimeraX^119,120^. To visualize AlphaFold2 model confidence score, both Pymol and ChimeraX were used. To calculate lipophilicity, pyMLP in ChimeraX was used. Helix angle measurements were performed using a Pymol plugin angle between helices.

### Electron microscopy

Third instar larvae and pupae brains at 6 hours after puparium formation (APF) were dissected in ice-cold 0.1M cacodylate buffer at pH 7.4 (EMS). The buffer was immediately replaced with a solution containing 2% glutaraldehyde, and 4% paraformaldehyde in 0.1M cacodylate buffer, and the brains were left to fix with gentle agitation for 30 minutes at room temperature before microwave-assisted fixation using a Pelco Biowave microwave (Ted Pella) with the following settings: 100W for 1 minute, followed by 1 minute OFF, repeated twice; then 450W for 20 seconds, followed by 20 seconds OFF, repeated five times. The brains were stored in the fixative overnight at 4°C before further processing.

Following the primary fixation, the brains underwent three 10-minute washes with fresh 0.1M sodium cacodylate buffer. Subsequently, the buffer was replaced with 2% osmium tetroxide (OsO_4_) diluted in 0.1M sodium cacodylate, 0.1M imidazole buffer, pH 7.5, and the samples were microwaved again at 100W for 1 minute, followed by 1 minute OFF (repeated twice), and then 450W for 20 seconds, with 20 seconds OFF, repeated five times. After this step, OsO_4_ was removed with three 10-minute washes using distilled water (ddH2O). En-bloc uranyl acetate (UA) staining was performed by adding saturated (8% w/v in ddH2O) UA to each tube and microwaving (450W for 1 minute, followed by 1 minute OFF, repeated once). The UA stain was then washed out with three 10-minute washes using ddH2O, before serial dehydration.

The samples were dehydrated using a series of ethanol (EtOH) dilutions at concentrations of 25%, 50%, 70%, 80%, and 95%, with each step followed by a microwave cycle at 250W for 45 seconds. Then, three changes of 100% EtOH were performed, each with a microwave cycle (250W for 1 minute, followed by 1 minute OFF, and then 250W for 1 minute). This was followed by three changes of 100% acetone, with the same microwave settings. The final acetone wash was used to transfer the samples to glass dram vials. The pure acetone was replaced with a 1:1 mixture of acetone and Embed 812 resin (EMS), and the samples were allowed to infiltrate overnight on a rotating shaker. All microwave steps were carried out with the samples in a chilled circulating water bath to maintain a temperature below 20°C.

The next day, the 1:1 acetone: Embed-812 mixture was replaced with 100% fresh Embed-812 resin and allowed to infiltrate for at least 1 hour on a rotating shaker at room temperature. Brains were then embedded in a thin layer of resin by flat-embedding between Aclar sheets and cured overnight in a 60°C oven. Embedded brains were examined on an Olympus upright microscope to check for any signs of damage. Intact, flat-embedded brains were then carefully trimmed near the region of interest (ROI) with a heated razor blade and re-embedded in Embed-812 using silicone flat embedding molds (EMS) to facilitate sectioning. 70nm sections were collected on 100-mesh formvar-coated grids and counterstained with 5% uranyl acetate for 20 minutes and Reynolds lead citrate for 7 minutes before being imaged using a Tecnai T12 electron microscope equipped with an AMT CCD camera and accompanying software

### Fluorescence microscopy and time-lapse imaging

A DeltaVision Elite Deconvolution Imaging System (GE Healthcare, Inc) equipped with a DIC imaging apparatus and a Photometrics CoolSnap HQ2 digital camera was used to capture fluorescence and DIC images. Applied Precision SoftWoRx 5.5 software was utilized for deconvolving and analyzing the images ^58^. To observe the engulfment process of C3 and the fusion between ventral hypodermal cells ABplaapppp and ABpraapppp by following the CED-1::GFP reporter, time-lapse recording was performed. Twelve serial Z-sections in 0.5 μm intervals between adjacent optical sections, from the ventral surface of to the middle of embryos, starting at 300 min-post the 1^st^ embryonic cell division, were collected using an established protocol ^58^. Images were captured every 2 min, with recordings typically lasting between 60 and 120 min. Embryos that exhibited normal elongation pattern were considered developing properly. The moment engulfment starts is defined as when the extension of pseudopods around C3 is first observed. The moment a nascent phagosome form is defined as when the pseudopods around a cell corpse join and make a full closure. The period between the budding and the sealing of the pseudopods is the time span of engulfment. The moment that ABplaapppp and ABpraapppp first touch is the time point when these two cells were 1^st^ seen to touch. If engulfed is not observed 30 min after the “0 min” time point, the apoptotic cell C3 is deemed “not engulfed”.

### RNAi treatment

*lpd-3* RNAi-by-feed construct was generated by inserting a 553 bp exon 15 of the *lpd-3* genomic DNA into pPD129.36 (a gift from Andrew Fire), the vector for the RNAi-by-feeding constructs. RNAi of *lpd-3* was performed by feeding ^121^, starting at mid-L4 stage. F1 embryos were collected for time-lapse recording 24 hours later. pPD129.36 was used as a negative control.

### Translating Ribosome Affinity Purification Sequencing

#### *Drosophila* strains

To generate flies in which ribosomes were tagged in astrocytes, we used alrm-Gal4, UAS-EGFP-L10a/+^48^ flies. wL3, 2 APF, and 6 APF animals were used and 150 CNSs were hand dissected in 1x PBS containing 100ng/mL of Cycloheximide for each sample.

#### Sample preparation

Dissected CNS were collected in an RNase free Eppendorf tube and kept on ice for no longer than 30min. Samples were then stored at -80°C.

#### Lysate preparation

Samples were combined to make a total of roughly 150 CNSs in each tube, Samples were homogenized in lysis buffer (20mM HEPES KOH pH7.4, 5mM MgCl_2_, 150mM KCl, protease inhibitor, with 0.5mM DTT, 100μg/mL cyclohexamide, 50μg/mL emetine, 10μl/mL SUPERase-In RNAse inhibitor and 10μl/mL RNase OUT ribonuclease inhibitor added right before use). 250μl buffer was added to each tube of 150 CNSs. Samples were then placed on ice and homogenized for 20-30s using a motorized hand-held homogenizer with pestles fitted to 1.5mL Eppendorf tube (Biomasher II, Kimble). After homogenization, lysis buffer was added to adult head samples to bring them up to 1mL. Samples were spun down at 2060g to remove nuclei. To each sample, 111μl NP-40 was added and was mixed gently by inversion. Then, 123μl DHPC was added, mixed gently by inversion and incubated on ice for 5min. Samples were centrifuged at 16,100g at 4°C for 15min.

#### Bead preparation

375μl of Protein G Dynalbeads were prepared for each sample. Beads were washed 3 times for 5min with shaking in a 0.15M KCl wash buffer (20mM HEPES-KOH pH7.4, 150mM KCl, 5mM MgCl_2_, 1% NP- 40, with 0.5mM DTT, 100μg/mL cyclohexamide and 50μg/mL emetine added immediately before use) and were pelleted using a magnetic stand. Stock solutions of cyclohexamide were prepared in MeOH at a concentration of 100mg/mL and emetine prepared in EtOH at a concentration of 50mg/mL. After the third wash, beads were resuspended in 0.15M KCl and GFP antibodies (50μg of each of HTZGFP-19C8 and HTZGFP-19F7) up to a total volume of 375μl per sample. Beads were incubated at RT for 2 hours with end-over-end rotation. Unbound antibody was removed by washing 3 times for 5min in the 0.15M KCl wash buffer.

#### Pull-down

For each sample, 1mL supernatant was transferred to the washed, GFP-bound beads. Beads were resuspended in sample lysates and incubated at 4°C with end over end rotation for 30min. Beads were then washed 3 times for 5min at 4°C in 0.35M KCl wash buffer (20mM HEPES-KOH pH7.4, 350mM KCl, 5mM MgCl_2_, 1%NP-40, with 0.5mM DTT, 100μg/mL cyclohexamide and 50μg/mL emetine added immediately before use). Samples were then placed on ice for up to 20min.

#### RNA recovery

Buffer was removed and samples were resuspended in RLT buffer (Qiagen RNA minielute kit) with BMe added just before use. Tubes was mixed by inversion to resuspend the beads and placed on ice. RNA isolation was performed according to the instructions provided in the kit. RNA was eluted by adding 10μl of water to the membrane, spinning for 1min, and then adding another 10μl of water and repeating the spin step.

#### RNA sequencing and analysis

RNA quality and quantity was assessed using an Agilent 2100 Bioanalyzer and RNA 60000 PicoChip, run with the eukaryotic total RNA program. Concentrations ranged from 5.1 ng-16.1 ng/ul as measured with the Aligent RNA 6000 Pico Assay. RIN values ranged from 6.4-7.3. cDNAs were prepared with SmarSeq Ultra Low input kit (Takara).

### Quantitative real-time PCR and Analysis

qPCR was performed on a QuantStudio 6 Flex using the SYBR Green Master mix (Roche). In all cases, samples were run simultaneously with three independent biological replicates for each target gene, and *rp49* was used as the reference gene (primers are in Table S1). To calculate changes in relative expression, using the 2^-ΔΔ^*^CT^* method^122^.

### Immunohistochemistry

Third instar larvae CNS were collected in 0.1% Triton X-100 (0.1% PBST) on ice. For samples collected at 2 and 6 hours after puparium formation (APF), white prepupae were collected and kept on a 1% agarose plate in a 25°C incubator for the required duration before dissection. In the case of samples collected at head eversion (HE), prepupae were placed on the 1% agarose plate, with the CNS of animals undergoing head eversion being dissected every 10 minutes. All dissected samples were placed in PBS with 0.1% Triton X-100 (0.1% PBST) on ice. Following this, the samples were transferred to a solution of 4% PFA in PBS and fixed for 20 minutes with gentle agitation at room temperature (RT). In most experiments, post-fixation, the samples were washed twice in 0.1% PBST, after which they were transferred to 0.1% PBST containing primary antibodies and incubated on a shaker at 4°C for 2 days. Three five-minute washes were performed at RT using 0.1% PBST, and then the samples were placed on a shaker in 0.1% PBST overnight at 4°C. Subsequently, the samples were moved to 0.1% PBST containing secondary antibodies for overnight incubation at 4°C, followed by further washing and storage in 0.1% PBST for a duration of up to 3 nights at 4°C. For α-Crz, the primary antibody incubation was conducted overnight, while the secondary antibody incubation lasted for 3 hours at room temperature. The washing steps were consistent across these procedures. Finally, the samples were mounted using Vectashield antifade reagent (Vector Laboratories). The primary antibodies used for staining larvae were as follows: α-Crz (Crz 1:1000, this study), α-GFP (abcam 1:1000), α-GAT (1:2000), and α-mCherry (Thermo Fisher).

### Western Blot

The samples were prepared according to the immunohistochemistry protocol. Each sample contained 10 CNSs, which were dissected and carefully transferred into PBS with Laemmeli Buffer containing β-Mercaptoethanol. The samples were then homogenized and boiled for 10 minutes. The samples were either frozen at -80°C or loaded directly into a 4-20% Tris-Bis Gel (Bio Rad). Gel electrophoresis was conducted at 100V for 1 hour to 1.5 hours in a Tris-glycine running buffer containing 8% SDS. The protein was then transferred onto a PVDF membrane in Tris-glycine buffer with 12% MeOH at 100V for 1 hour or overnight at 33V.

The membrane was blocked in 5% BSA in 0.1% TBST for 1 hour at room temperature and subsequently incubated in the primary antibody overnight at 4°C. Afterward, the blot underwent three washes for 15 minutes each in 0.1% TBST and was then incubated overnight in HRP-conjugated secondary antibodies (1:5000, Jackson Immuno Research) in blocking solution at 4°C. Following additional washing steps as described, the blots were incubated in ECL (Bio Rad). Blots were briefly washed in PBS and imaged on a Chemidoc MP Imaging System (Bio Rad). Optical density was quantified using Image J’s Gel Analyzer tool. The primary antibodies used for Western blots were GFP (1:1000 Cell Signaling 29456), Tubulin (1:1000 Developmental Studies Hybridoma Bank E7), Actin (1:1000 Cell Signaling) and Brp (1:1000 DSHB).

### Image acquisition and processing

Fixed fly CNS samples were imaged using a Zeiss LSM 880 with Airyscan confocal microscope. Live imaging was conducted using either the Zeiss LSM 880 with Airyscan confocal microscope or the Zeiss Axio Examiner equipped with a Yokogawa spinning disk and Hamamatsu camera. High-resolution astrocyte imaging involved acquiring confocal z-stacks with optimal z-intervals and approximately 0.05-0.09 µm/pixel resolution, utilizing a 40x/1.3 Plan-Apochromat oil objective. In cases where needed, image tiles were stitched together, and all images were processed using Airyscan.

## QUANTIFICATION AND STATISTICAL ANALYSIS

The investigator performing the image analysis was blinded to genotype and condition. To quantify vCrz debris the VNC was outlined and the area of vCrz debris was collected. All the images for the experiments used the same threshold and the area of all nuclei was eliminated. The relative percent area to larval control was used for analysis. To quantify vesicles 4 independent regions of a VNC or a clone were selected and vesicle size and numbers were manually counted. To quantify ER-PM contact numbers, 4 independent regions of the VNC in identical region was selected. Using ImageJ background was corrected and using a consistent threshold the ER-PM contacts sites were counted. All statistical analyses were carried out using GraphPad Prism 8/9; statistical details, p values, and numbers of analyzed samples are indicated in the figure legends.

## SUPPLEMENTAL ITEMS

**Movie S1**. Live imaging of Crz neuron remodeling was conducted using GFP-labeled Crz neurons (*Crz-Gal4, UAS-mCD8-GFP*) in an *ex vivo* preparation. Drift correction and adjustment of GFP intensity were performed using Image J.

**Movie S2**. Three independent events of early phagosome fusion were imaged using *ex vivo* live imaging preparations. Early phagosomes were labeled with 2xGF-FYVE (*GMR25h07-Gal4, UAS-2x GFP-FYVE*), and drift correction and GFP intensity adjustments were carried out using Image J.

**Movie S3**. Movie showing the maturation of early phagosomes into phagolysosomes (*GMR25h07-Gal4, UAS- 2x GFP-FYVE, UAS-Spinster-mRFP*). Drift correction and adjustment of GFP intensity were performed using Image J.

**Supplementary Table 1.** Results of the screen to identify glial genes that regulate Crz neuron remodeling.

**Supplementary Table 2.** Comparison of genes known to be critical in axonal debris phagocytosis during Wallerian degeneration versus vCrz phagocytosis during development.

## Notes

### Competing Interest Statement

The authors have declared no competing interest.

## References

1. Landmesser, L., and Pilar, G. (1972). The onset and development of transmission in the chick ciliary ganglion. J Physiol 222, 691–713. 10.1113/jphysiol.1972.sp009822.

2. Landmesser, L., and Pilar, G. (1974). Synaptic transmission and cell death during normal ganglionic development. J Physiol 241, 737–749. 10.1113/jphysiol.1974.sp010681.

3. Southwell, D.G., Paredes, M.F., Galvao, R.P., Jones, D.L., Froemke, R.C., Sebe, J.Y., Alfaro-Cervello, C., Tang, Y., Garcia-Verdugo, J.M., Rubenstein, J.L., et al. (2012). Intrinsically determined cell death of developing cortical interneurons. Nature 491, 109–113. 10.1038/NATURE11523.

4. Faust, T.E., Gunner, G., and Schafer, D.P. (2021). Mechanisms governing activity-dependent synaptic pruning in the mammalian CNS. Nat Rev Neurosci 22, 657. 10.1038/S41583-021-00507-Y.

5. Riccomagno, M.M., and Kolodkin, A.L. (2015). Sculpting neural circuits by axon and dendrite pruning. Annu Rev Cell Dev Biol 31, 779–805. 10.1146/ANNUREV-CELLBIO-100913-013038.

6. Neukomm, L.J., and Freeman, M.R. (2014). Diverse cellular and molecular modes of axon degeneration. Trends Cell Biol 24, 515–523. 10.1016/J.TCB.2014.04.003.

7. Chen, C., and Regehr, W.G. (2000). Developmental remodeling of the retinogeniculate synapse. Neuron 28, 955–966. 10.1016/S0896-6273(00)00166-5.

8. Cuadros, M.A., and Navascués, J. (1998). The origin and differentiation of microglial cells during development. Prog Neurobiol 56, 173–189. 10.1016/S0301-0082(98)00035-5.

9. Parnaik, R., Raff, M.C., and Scholes, J. (2000). Differences between the clearance of apoptotic cells by professional and non-professional phagocytes. Curr Biol 10, 857–860. 10.1016/S0960-9822(00)00598-4.

10. Lee, J.H., Kim, J. young Noh, S., Lee, H., Lee, S.Y., Mun, J.Y., Park, H., and Chung, W.S. (2021). Astrocytes phagocytose adult hippocampal synapses for circuit homeostasis. Nature 590, 612–617. 10.1038/s41586-020-03060-3.

11. Auguste, Y.S.S., Ferro, A., Kahng, J.A., Xavier, A.M., Dixon, J.R., Vrudhula, U., Nichitiu, A.S., Rosado, D., Wee, T.L., Pedmale, U. V., et al. (2022). Oligodendrocyte precursor cells engulf synapses during circuit remodeling in mice. Nat Neurosci 25, 1273–1278. 10.1038/S41593-022-01170-X.

12. Buchanan, J.A., Elabbady, L., Collman, F., Jorstad, N.L., Bakken, T.E., Ott, C., Glatzer, J., Bleckert, A.A., Bodor, A.L., Brittain, D., et al. (2022). Oligodendrocyte precursor cells ingest axons in the mouse neocortex. Proc Natl Acad Sci U S A 119, e2202580119. 10.1073/PNAS.2202580119/SUPPL_FILE/PNAS.2202580119.SM04.MOV.

13. Ashwell, K. (1990). Microglia and cell death in the developing mouse cerebellum. Developmental Brain Research 55, 219–230. 10.1016/0165-3806(90)90203-B.

14. Marínmarímarín-Teva, J.L., Cuadros, M.A., Calvente, R., Almendros, A., and Navascué, J. (1999). Naturally Occurring Cell Death and Migration of Microglial Precursors in the Quail Retina During Normal Development. J. Comp. Neurol 412, 255–275. 10.1002/(SICI)1096-9861(19990920)412:2.

15. Marín-Teva, J.L., Dusart, I., Colin, C., Gervais, A., Van Rooijen, N., and Mallat, M. (2004). Microglia Promote the Death of Developing Purkinje Cells. Neuron 41, 535–547. 10.1016/S0896-6273(04)00069-8.

16. Hong, S., Beja-Glasser, V.F., Nfonoyim, B.M., Frouin, A., Li, S., Ramakrishnan, S., Merry, K.M., Shi, Q., Rosenthal, A., Barres, B.A., et al. (2016). Complement and microglia mediate early synapse loss in Alzheimer mouse models. Science 352, 712–716. 10.1126/SCIENCE.AAD8373.

17. Tasdemir-Yilmaz, O.E., and Freeman, M.R. (2014). Astrocytes engage unique molecular programs to engulf pruned neuronal debris from distinct subsets of neurons. Genes Dev 28, 20–33. 10.1101/gad.229518.113.

18. Chung, W.S., Clarke, L.E., Wang, G.X., Stafford, B.K., Sher, A., Chakraborty, C., Joung, J., Foo, L.C., Thompson, A., Chen, C., et al. (2013). Astrocytes mediate synapse elimination through MEGF10 and MERTK pathways. Nature 504, 394–400. 10.1038/NATURE12776.

19. Stevens, B., Allen, N.J., Vazquez, L.E., Howell, G.R., Christopherson, K.S., Nouri, N., Micheva, K.D., Mehalow, A.K., Huberman, A.D., Stafford, B., et al. (2007). The Classical Complement Cascade Mediates CNS Synapse Elimination. Cell 131, 1164–1178. 10.1016/j.cell.2007.10.036.

20. Schafer, D.P., Lehrman, E.K., Kautzman, A.G., Koyama, R., Mardinly, A.R., Yamasaki, R., Ransohoff, R.M., Greenberg, M.E., Barres, B.A., and Stevens, B. (2012). Microglia Sculpt Postnatal Neural Circuits in an Activity and Complement-Dependent Manner. Neuron 74, 691. 10.1016/J.NEURON.2012.03.026.

21. Gunner, G., Cheadle, L., Johnson, K.M., Ayata, P., Badimon, A., Mondo, E., Nagy, M.A., Liu, L., Bemiller, S.M., Kim, K.W., et al. (2019). Sensory lesioning induces microglial synapse elimination via ADAM10 and fractalkine signaling. Nat Neurosci 22, 1075–1088. 10.1038/s41593-019-0419-y.

22. Filipello, F., Morini, R., Corradini, I., Zerbi, V., Canzi, A., Michalski, B., Erreni, M., Markicevic, M., Starvaggi-Cucuzza, C., Otero, K., et al. (2018). The Microglial Innate Immune Receptor TREM2 Is Required for Synapse Elimination and Normal Brain Connectivity. Immunity 48, 979–991.e8. 10.1016/J.IMMUNI.2018.04.016.

23. Jay, T.R., von Saucken, V.E., Muñoz, B., Codocedo, J.F., Atwood, B.K., Lamb, B.T., and Landreth, G.E. (2019). TREM2 is required for microglial instruction of astrocytic synaptic engulfment in neurodevelopment. Glia 67, 1873–1892. 10.1002/GLIA.23664.

24. Podleśny-Drabiniok, A., Marcora, E., and Goate, A.M. (2020). Microglial Phagocytosis: A Disease-Associated Process Emerging from Alzheimer’s Disease Genetics. Trends Neurosci 43, 965. 10.1016/J.TINS.2020.10.002.

25. Sierra, A., Abiega, O., Shahraz, A., and Neumann, H. (2013). Janus-faced microglia: beneficial and detrimental consequences of microglial phagocytosis. Front Cell Neurosci 7. 10.3389/FNCEL.2013.00006.

26. Levin, R., Grinstein, S., and Canton, J. (2016). The life cycle of phagosomes: formation, maturation, and resolution.

27. Lee, H.J., Woo, Y., Hahn, T.W., Jung, Y.M., and Jung, Y.J. (2020). Formation and Maturation of the Phagosome: A Key Mechanism in Innate Immunity against Intracellular Bacterial Infection. Microorganisms 8, 1–22. 10.3390/MICROORGANISMS8091298.

28. Kinchen, J.M., and Ravichandran, K.S. (2008). Phagosome maturation: Going through the acid test. Preprint, 10.1038/nrm2515 10.1038/nrm2515.

29. Hakim, Y., Yaniv, S.P., and Schuldiner, O. (2014). Astrocytes play a key role in Drosophila mushroom body axon pruning. PLoS One 9. 10.1371/journal.pone.0086178.

30. Choi, Y.J., Lee, G., and Park, J.H. (2006). Programmed cell death mechanisms of identifiable peptidergic neurons in drosophila melanogaster. Development 133, 2223–2232. 10.1242/dev.02376.

31. Lee, G., Wang, Z., Sehgal, R., Chen, C.H., Kikuno, K., Hay, B., and Park, J.H. (2011). Drosophila caspases involved in developmentally regulated programmed cell death of peptidergic neurons during early metamorphosis. Journal of Comparative Neurology 519, 34–48. 10.1002/cne.22498.

32. Wu, Y.C., and Horvitz, H.R. (1998). The C. elegans cell corpse engulfment gene ced-7 encodes a protein similar to ABC transporters. Cell 93, 951–960. 10.1016/S0092-8674(00)81201-5.

33. Morizawa, Y.M., Hirayama, Y., Ohno, N., Shibata, S., Shigetomi, E., Sui, Y., Nabekura, J., Sato, K., Okajima, F., Takebayashi, H., et al. (2017). Reactive astrocytes function as phagocytes after brain ischemia via ABCA1-mediated pathway. Nat Commun 8. 10.1038/s41467-017-00037-1.

34. Awasaki, T., Huang, Y., O’Connor, M.B., and Lee, T. (2011). Glia instruct developmental neuronal remodeling through TGF-Î 2 signaling. Nat Neurosci 14, 821–823. 10.1038/nn.2833.

35. Lu, T.Y., Doherty, J., and Freeman, M.R. (2014). DRK/DOS/SOS converge with Crk/Mbc/dCed-12 to activate Rac1 during glial engulfment of axonal debris. Proc Natl Acad Sci U S A 111, 12544–12549. 10.1073/pnas.1403450111.

36. Ziegenfuss, J.S., Doherty, J., and Freeman, M.R. (2012). Distinct molecular pathways mediate glial activation and engulfment of axonal debris after axotomy. Nat Neurosci 15, 979–987. 10.1038/nn.3135.

37. MacDonald, J.M., Doherty, J., Hackett, R., and Freeman, M.R. (2013). The c-Jun kinase signaling cascade promotes glial engulfment activity through activation of draper and phagocytic function. Cell Death Differ 20, 1140–1148. 10.1038/cdd.2013.30.

38. Logan, M.A., Hackett, R., Doherty, J., Sheehan, A., Speese, S.D., and Freeman, M.R. (2012). Negative regulation of glial engulfment activity by Draper terminates glial responses to axon injury. Nat Neurosci 15, 722–730. 10.1038/nn.3066.

39. Doherty, J., Sheehan, A.E., Bradshaw, R., Fox, A.N., Lu, T.Y., and Freeman, M.R. (2014). PI3K Signaling and Stat92E Converge to Modulate Glial Responsiveness to Axonal Injury. PLoS Biol 12. 10.1371/journal.pbio.1001985.

40. MacDonald, J.M., Beach, M.G., Porpiglia, E., Sheehan, A.E., Watts, R.J., and Freeman, M.R. (2006). The Drosophila Cell Corpse Engulfment Receptor Draper Mediates Glial Clearance of Severed Axons. Neuron 50, 869–881. 10.1016/j.neuron.2006.04.028.

41. Lu, T.Y., MacDonald, J.M., Neukomm, L.J., Sheehan, A.E., Bradshaw, R., Logan, M.A., and Freeman, M.R. (2017). Axon degeneration induces glial responses through Draper-TRAF4-JNK signalling. Nat Commun 8. 10.1038/ncomms14355.

42. Ziegenfuss, J.S., Biswas, R., Avery, M.A., Hong, K., Sheehan, A.E., Yeung, Y.-G., Stanley, E.R., and Freeman, M.R. Draper-dependent glial phagocytic activity is mediated by Src and Syk family kinase signalling.

43. Verstreken, P., Ohyama, T., Haueter, C., Habets, R.L.P., Lin, Y.Q., Swan, L.E., Ly, C. V., Venken, K.J.T., De Camilli, P., and Bellen, H.J. (2009). Tweek, an Evolutionarily Conserved Protein, Is Required for Synaptic Vesicle Recycling. Neuron 63, 203–215. 10.1016/j.neuron.2009.06.017.

44. Wang, C., Wang, B., Pandey, T., Long, Y., Zhang, J., Oh, F., Sima, J., Guo, R., Liu, Y., Zhang, C., et al. (2022). A conserved megaprotein-based molecular bridge critical for lipid trafficking and cold resilience. Nature Communications 2022 13:1 *13*, 1–11. 10.1038/s41467-022-34450-y.

45. Doherty, J., Logan, M.A., Taşdemir, Ö.E., and Freeman, M.R. (2009). Ensheathing glia function as phagocytes in the adult Drosophila brain. Journal of Neuroscience 29, 4768–4781. 10.1523/JNEUROSCI.5951-08.2009.

46. Muthukumar, A.K., Stork, T., and Freeman, M.R. (2014). Activity-dependent regulation of astrocyte GAT levels during synaptogenesis. Nat Neurosci 17, 1340–1350. 10.1038/nn.3791.

47. Wagh, D.A., Rasse, T.M., Asan, E., Hofbauer, A., Schwenkert, I., Dürrbeck, H., Buchner, S., Dabauvalle, M.C., Schmidt, M., Qin, G., et al. (2006). Bruchpilot, a protein with homology to ELKS/CAST, is required for structural integrity and function of synaptic active zones in Drosophila. Neuron 49, 833–844. 10.1016/J.NEURON.2006.02.008.

48. Huang, Y., Ng, F.S., and Jackson, F.R. (2015). Comparison of larval and adult Drosophila astrocytes reveals stage-specific gene expression profiles. G3 (Bethesda) 5, 551–558. 10.1534/G3.114.016162.

49. Wucherpfennig, T., Wilsch-Bräuninger, M., and González-Gaitán, M. (2003). Role of Drosophila Rab5 during endosomal trafficking at the synapse and evoked neurotransmitter release. Journal of Cell Biology 161, 609–624. 10.1083/JCB.200211087.

50. Li, T., Fu, T.M., Wong, K.K.L., Li, H., Xie, Q., Luginbuhl, D.J., Wagner, M.J., Betzig, E., and Luo, L. (2021). Cellular bases of olfactory circuit assembly revealed by systematic time-lapse imaging. Cell 184, 5107–5121.e14. 10.1016/J.CELL.2021.08.030.

51. He, F., Agosto, M.A., Nichols, R.M., Kailasam, L., and Wensel, T.G. (2018). Vps34 and PI(3)P are critical for autophagy, phagocytosis, and endosome processing in RPE cells. Invest Ophthalmol Vis Sci 59, 4016–4016.

52. Naufer, A., Hipolito, V.E.B., Ganesan, S., Prashar, A., Zaremberg, V., Botelho, R.J., and Terebiznik, M.R. (2018). pH of endophagosomes controls association of their membranes with Vps34 and PtdIns(3)P levels. Journal of Cell Biology 217, 329–346. 10.1083/jcb.201702179.

53. Sweeney, S., Neuron, G.D.-, and 2002, undefined (2002). Unrestricted synaptic growth in spinster—a late endosomal protein implicated in TGF-β-mediated synaptic growth regulation. cell.comST Sweeney, GW DavisNeuron, 2002•cell.com 36, 403–416.

54. Nakano, Y., Fujitani, K., Kurihara, J., Ragan, J., Usui-Aoki, K., Shimoda, L., Lukacsovich, T., Suzuki, K., Sezaki, M., Sano, Y., et al. (2001). Mutations in the Novel Membrane Protein Spinster Interfere with Programmed Cell Death and Cause Neural Degeneration in Drosophila melanogaster. Mol Cell Biol 21, 3775–3788. 10.1128/MCB.21.11.3775-3788.2001.

55. Dermaut, B., Norga, K.K., Kania, A., Verstreken, P., Pan, H., Zhou, Y., Callaerts, P., and Bellen, H.J. (2005). Aberrant lysosomal carbohydrate storage accompanies endocytic defects and neurodegeneration in Drosophila benchwarmer. J Cell Biol 170, 127. 10.1083/JCB.200412001.

56. Lin, H.J., Herman, P., Kang, J.S., and Lakowicz, J.R. (2001). Fluorescence lifetime characterization of novel low-pH probes. Anal Biochem 294, 118–125. 10.1006/ABIO.2001.5155.

57. Zhou, Z., Hartwieg, E., and Horvitz, H.R. (2001). CED-1 is a transmembrane receptor that mediates cell corpse engulfment in C. elegans. Cell 104, 43–56. 10.1016/S0092-8674(01)00190-8.

58. Lu, N., Yu, X., He, X., and Zhou, Z. (2009). Detecting apoptotic cells and monitoring their clearance in the nematode Caenorhabditis elegans. Methods Mol Biol 559, 357–370. 10.1007/978-1-60327-017-5_25.

59. Jumper, J., Evans, R., Pritzel, A., Green, T., Figurnov, M., Ronneberger, O., Tunyasuvunakool, K., Bates, R., Žídek, A., Potapenko, A., et al. (2021). Highly accurate protein structure prediction with AlphaFold. Nature 2021 596:7873 596, 583–589. 10.1038/s41586-021-03819-2.

60. Toulmay, A., Whittle, F.B., Yang, J., Bai, X., Diarra, J., Banerjee, S., Levine, T.P., Golden, A., and Prinz, W.A. (2022). Vps13-like proteins provide phosphatidylethanolamine for GPI anchor synthesis in the ER. Journal of Cell Biology 221. 10.1083/jcb.202111095.

61. Braschi, B., Bruford, E.A., Cavanagh, A.T., Neuman, S.D., and Bashirullah, A. (2022). The bridge-like lipid transfer protein (BLTP) gene group: introducing new nomenclature based on structural homology indicating shared function. Hum Genomics 16. 10.1186/s40246-022-00439-3.

62. Neuman, S.D., Levine, T.P., and Bashirullah, A. (2022). A novel superfamily of bridge-like lipid transfer proteins. Trends Cell Biol 32, 962–974. 10.1016/J.TCB.2022.03.011.

63. Neuman, S.D., Jorgensen, J.R., Cavanagh, A.T., Smyth, J.T., Selegue, J.E., Emr, S.D., and Bashirullah, A. (2022). The Hob proteins are novel and conserved lipid-binding proteins at ER-PM contact sites. J Cell Sci 135. 10.1242/JCS.259086/VIDEO-2.

64. Manford, A.G., Stefan, C.J., Yuan, H.L., MacGurn, J.A., and Emr, S.D. (2012). ER-to-plasma membrane tethering proteins regulate cell signaling and ER morphology. Dev Cell 23, 1129–1140. 10.1016/J.DEVCEL.2012.11.004.

65. Ugrankar, R., Bowerman, J., Hariri, H., Chandra, M., Chen, K., Bossanyi, M.F., Datta, S., Rogers, S., Eckert, K.M., Vale, G., et al. (2019). Drosophila Snazarus Regulates a Lipid Droplet Population at Plasma Membrane-Droplet Contacts in Adipocytes. Dev Cell 50, 557–572.e5. 10.1016/J.DEVCEL.2019.07.021.

66. Chang, C.L., Hsieh, T.S., Yang, T.T., Rothberg, K.G., Azizoglu, D.B., Volk, E., Liao, J.C., and Liou, J. (2013). Feedback regulation of receptor-induced Ca2+ signaling mediated by E-Syt1 and Nir2 at endoplasmic reticulum-plasma membrane junctions. Cell Rep 5, 813–825. 10.1016/J.CELREP.2013.09.038.

67. Cheng, S., Wang, K., Zou, W., Miao, R., Huang, Y., Wang, H., and Wang, X. (2015). PtdIns(4,5)P2 and PtdIns3P coordinate to regulate phagosomal sealing for apoptotic cell clearance. Journal of Cell Biology 210, 485–502. 10.1083/JCB.201501038/VIDEO-9.

68. Bohdanowicz, M., and Grinstein, S. (2013). Role of phospholipids in endocytosis, phagocytosis, and macropinocytosis. Physiol Rev 93, 69–106. 10.1152/PHYSREV.00002.2012/ASSET/IMAGES/LARGE/Z9J0011326370004.JPEG.

69. Di Paolo, G., and De Camilli, P. (2006). Phosphoinositides in cell regulation and membrane dynamics. Nature 2006 443:7112 443, 651–657. 10.1038/nature05185.

70. Várnai, P., and Balla, T. (1998). Visualization of phosphoinositides that bind pleckstrin homology domains: calcium-and agonist-induced dynamic changes and relationship to myo-[3H]inositol-labeled phosphoinositide pools. J Cell Biol 143, 501–510. 10.1083/JCB.143.2.501.

71. Jay, T.R., Kang, Y., Jefferson, A., and Freeman, M.R. (2021). An ELISA-based method for rapid genetic screens in Drosophila. Proc Natl Acad Sci U S A 118. 10.1073/PNAS.2107427118.

72. Gueneau, L., Fish, R.J., Shamseldin, H.E., Voisin, N., Tran Mau-Them, F., Preiksaitiene, E., Monroe, G.R., Lai, A., Putoux, A., Allias, F., et al. (2018). KIAA1109 Variants Are Associated with a Severe Disorder of Brain Development and Arthrogryposis. Am J Hum Genet 102, 116–132. 10.1016/J.AJHG.2017.12.002.

73. Kumar, K., Bellad, A., Bellad, A., Prasad, P., Girimaji, S.C., Muthusamy, B., and Muthusamy, B. (2020). KIAA1109 gene mutation in surviving patients with Alkuraya-Kučinskas syndrome: A review of literature. BMC Med Genet 21, 1–11. 10.1186/s12881-020-01074-2.

74. Kane, M.S., Diamonstein, C.J., Hauser, N., Deeken, J.F., Niederhuber, J.E., and Vilboux, T. (2019). Endosomal trafficking defects in patient cells with KIAA1109 biallelic variants. Genes Dis 6, 56–67. 10.1016/J.GENDIS.2018.12.004.

75. Hakim, Y., Yaniv, S.P., and Schuldiner, O. (2014). Astrocytes play a key role in Drosophila mushroom body axon pruning. PLoS One 9. 10.1371/JOURNAL.PONE.0086178.

76. Wang, Z., Lee, G., Vuong, R., and Park, J.H. (2019). Two-factor specification of apoptosis: TGF-β signaling acts cooperatively with ecdysone signaling to induce cell-and stage-specific apoptosis of larval neurons during metamorphosis in Drosophila melanogaster. Apoptosis 24, 972–989. 10.1007/s10495-019-01574-4.

77. Haney, M.S., Bohlen, C.J., Morgens, D.W., Ousey, J.A., Barkal, A.A., Tsui, C.K., Ego, B.K., Levin, R., Kamber, R.A., Collins, H., et al. (2018). Identification of phagocytosis regulators using magnetic genome-wide CRISPR screens. Nat Genet 50, 1716–1727. 10.1038/s41588-018-0254-1.

78. Kinchen, J.M., Doukoumetzidis, K., Almendinger, J., Stergiou, L., Tosello-Trampont, A., Sifri, C.D., Hengartner, M.O., and Ravichandran, K.S. (2008). A pathway for phagosome maturation during engulfment of apoptotic cells. Nat Cell Biol 10, 556–566. 10.1038/ncb1718.

79. Jeng, E.E., Bhadkamkar, V., Ibe, N.U., Gause, H., Jiang, L., Chan, J., Jian, R., Jimenez-Morales, D., Stevenson, E., Krogan, N.J., et al. (2019). Systematic Identification of Host Cell Regulators of Legionella pneumophila Pathogenesis Using a Genome-wide CRISPR Screen. Cell Host Microbe 26, 551–563.e6. 10.1016/J.CHOM.2019.08.017.

80. Hanna, M., Guillén-Samander, A., and Camilli, P. De (2023). RBG Motif Bridge-Like Lipid Transport Proteins: Structure, Functions, and Open Questions. 10.1146/annurev-cellbio-120420-014634 39. 10.1146/ANNUREV-CELLBIO-120420-014634.

81. Kumar, N., Leonzino, M., Hancock-Cerutti, W., Horenkamp, F.A., Li, P.Q., Lees, J.A., Wheeler, H., Reinisch, K.M., and De Camilli, P. (2018). VPS13A and VPS13C are lipid transport proteins differentially localized at ER contact sites. Journal of Cell Biology 217, 3625–3639. 10.1083/JCB.201807019.

82. Zaman, M.F., Nenadic, A., Radojičić, A., Rosado, A., and Beh, C.T. (2020). Sticking With It: ER-PM Membrane Contact Sites as a Coordinating Nexus for Regulating Lipids and Proteins at the Cell Cortex. Preprint at Frontiers Media S.A., 10.3389/fcell.2020.00675 10.3389/fcell.2020.00675.

83. Leonzino, M., Reinisch, K.M., and De Camilli, P. (2021). Insights into VPS13 properties and function reveal a new mechanism of eukaryotic lipid transport. Biochim Biophys Acta Mol Cell Biol Lipids 1866. 10.1016/j.bbalip.2021.159003.

84. Dabrowski, R., Tulli, S., and Graef, M. (2023). Parallel phospholipid transfer by Vps13 and Atg2 determines autophagosome biogenesis dynamics. J Cell Biol 222. 10.1083/jcb.202211039.

85. Gómez-Sánchez, R., Rose, J., Guimarães, R., Mari, M., Papinski, D., Rieter, E., Geerts, W.J., Hardenberg, R., Kraft, C., Ungermann, C., et al. (2018). Atg9 establishes Atg2-dependent contact sites between the endoplasmic reticulum and phagophores. Journal of Cell Biology 217, 2743–2763. 10.1083/JCB.201710116/VIDEO-5.

86. Osawa, T., and Noda, N.N. (2019). Atg2: A novel phospholipid transfer protein that mediates de novo autophagosome biogenesis. Protein Sci 28, 1005. 10.1002/PRO.3623.

87. Valverde, D.P., Yu, S., Boggavarapu, V., Kumar, N., Lees, J.A., Walz, T., Reinisch, K.M., and Melia, T.J. (2019). ATG2 transports lipids to promote autophagosome biogenesis. Journal of Cell Biology 218, 1787–1798. 10.1083/JCB.201811139.

88. Osawa, T., Kotani, T., Kawaoka, T., Hirata, E., Suzuki, K., Nakatogawa, H., Ohsumi, Y., and Noda, N.N. (2019). Atg2 mediates direct lipid transfer between membranes for autophagosome formation. Nature Structural & Molecular Biology 2019 26:4 26, 281–288. 10.1038/s41594-019-0203-4.

89. Park, J.S., and Neiman, A.M. (2012). VPS13 regulates membrane morphogenesis during sporulation in Saccharomyces cerevisiae. J Cell Sci 125, 3004–3011. 10.1242/JCS.105114/-/DC1.

90. Nakamura, T.S., Suda, Y., Muneshige, K., Fujieda, Y., Okumura, Y., Inoue, I., Tanaka, T., Takahashi, T., Nakanishi, H., Gao, X.D., et al. (2021). Suppression of Vps13 adaptor protein mutants reveals a central role for PI4P in regulating prospore membrane extension. PLoS Genet 17. 10.1371/JOURNAL.PGEN.1009727.

91. DP, V., S, Y., V, B., N, K., JA, L., T, W., KM, R., and TJ, M. (2019). ATG2 transports lipids to promote autophagosome biogenesis. J Cell Biol 218, 1787–1798. 10.1083/JCB.201811139.

92. Wang, C., Wang, B., Pandey, T., Long, Y., Zhang, J., Sima, J., Guo, R., Liu, Y., Zhang, C., Mukherjee, S., et al. A megaprotein-based molecular bridge critical for lipid trafficking and cold resilience. 10.1101/2022.08.25.505359.

93. Park, J.S., Okumura, Y., Tachikawa, H., and Neiman, A.M. (2013). SPO71 encodes a developmental stage-specific partner for Vps13 in Saccharomyces cerevisiae. Eukaryot Cell 12, 1530–1537. 10.1128/EC.00239-13/ASSET/CEB05665-39C9-4898-99CA-96AB8823B92A/ASSETS/GRAPHIC/ZEK9990941840006.JPEG.

94. Adlakha, J., Hong, Z., Li, P., and Reinisch, K.M. (2022). Structural and biochemical insights into lipid transport by VPS13 proteins. J Cell Biol 221. 10.1083/JCB.202202030.

95. Peter, A.T.J., Herrmann, B., Antunes, D., Rapaport, D., Dimmer, K.S., and Kornmann, B. (2017). Vps13- Mcp1 interact at vacuole–mitochondria interfaces and bypass ER–mitochondria contact sites. Journal of Cell Biology 216, 3219–3229. 10.1083/JCB.201610055.

96. Bean, B.D.M., Dziurdzik, S.K., Kolehmainen, K.L., Fowler, C.M.S., Kwong, W.K., Grad, L.I., Davey, M., Schluter, C., and Conibear, E. (2018). Competitive organelle-specific adaptors recruit Vps13 to membrane contact sites. Journal of Cell Biology 217, 3593–3607. 10.1083/JCB.201804111.

97. Maeda, S., Yamamoto, H., Kinch, L.N., Garza, C.M., Takahashi, S., Otomo, C., Grishin, N. V., Forli, S., Mizushima, N., and Otomo, T. (2020). Structure, lipid scrambling activity and role in autophagosome formation of ATG9A. Nature Structural & Molecular Biology 2020 27:12 27, 1194–1201. 10.1038/s41594-020-00520-2.

98. Matoba, K., Kotani, T., Tsutsumi, A., Tsuji, T., Mori, T., Noshiro, D., Sugita, Y., Nomura, N., Iwata, S., Ohsumi, Y., et al. (2020). Atg9 is a lipid scramblase that mediates autophagosomal membrane expansion. Nature Structural & Molecular Biology 2020 27:12 27, 1185–1193. 10.1038/s41594-020-00518-w.

99. Park, J.S., Hu, Y., Hollingsworth, N.M., Miltenberger-Miltenyi, G., and Neiman, A.M. (2022). Interaction between VPS13A and the XK scramblase is important for VPS13A function in humans. J Cell Sci 135. 10.1242/JCS.260227/276291/AM/INTERACTION-BETWEEN-VPS13A-AND-THE-XK-SCRAMBLASE.

100. Guillen-Samander, A., Wu, Y., Sebastian Pineda, S., Garcıa, F.J., Eisen, J.N., Leonzino, M., Ugur, B., Kellis, M., Heiman, M., and De Camilli, P. (2022). A partnership between the lipid scramblase XK and the lipid transfer protein VPS13A at the plasma membrane. Proc Natl Acad Sci U S A 119, e2205425119. 10.1073/PNAS.2205425119/SUPPL_FILE/PNAS.2205425119.SM02.MP4.

101. Ueno, S.I., Maruki, Y., Nakamura, M., Tomemori, Y., Kamae, K., Tanabe, H., Yamashita, Y., Matsuda, S., Kaneko, S., and Sano, A. (2001). The gene encoding a newly discovered protein, chorein, is mutated in chorea-acanthocytosis. Nature Genetics 2001 28:2 28, 121–122. 10.1038/88825.

102. Rampoldi, L., Dobson-Stone, C., Rubio, J.P., Danek, A., Chalmers, R.M., Wood, N.W., Verellen, C., Ferrer, X., Malandrini, A., Fabrizi, G.M., et al. (2001). A conserved sorting-associated protein is mutant in chorea-acanthocytosis. Nature Genetics 2001 28:2 28, 119–120. 10.1038/88821.

103. Kolehmainen, J., Black, G.C.M., Saarinen, A., Chandler, K., Clayton-Smith, J., Träskelin, A.L., Perveen, R., Kivitie-Kallio, S., Norio, R., Warburg, M., et al. (2003). Cohen syndrome is caused by mutations in a novel gene, COH1, encoding a transmembrane protein with a presumed role in vesicle-mediated sorting and intracellular protein transport. Am J Hum Genet 72, 1359–1369. 10.1086/375454.

104. Schormair, B., Kemlink, D., Mollenhauer, B., Fiala, O., Machetanz, G., Roth, J., Berutti, R., Strom, T.M., Haslinger, B., Trenkwalder, C., et al. (2018). Diagnostic exome sequencing in early-onset Parkinson’s disease confirms VPS13C as a rare cause of autosomal-recessive Parkinson’s disease. Clin Genet 93, 603–612. 10.1111/CGE.13124.

105. Lesage, S., Drouet, V., Majounie, E., Deramecourt, V., Jacoupy, M., Nicolas, A., Cormier-Dequaire, F., Hassoun, S.M., Pujol, C., Ciura, S., et al. (2016). Loss of VPS13C Function in Autosomal-Recessive Parkinsonism Causes Mitochondrial Dysfunction and Increases PINK1/Parkin-Dependent Mitophagy. Am J Hum Genet 98, 500–513. 10.1016/j.ajhg.2016.01.014.

106. Darvish, H., Bravo, P., Tafakhori, A., Azcona, L.J., Ranji-Burachaloo, S., Johari, A.H., and Paisán-Ruiz, C. (2018). Identification of a large homozygous VPS13C deletion in a patient with early-onset Parkinsonism. Movement Disorders 33, 1968–1970. 10.1002/MDS.27516.

107. Dziurdzik, S.K., Bean, B.D.M., Davey, M., and Conibear, E. (2020). A VPS13D spastic ataxia mutation disrupts the conserved adaptor-binding site in yeast Vps13. Hum Mol Genet 29, 635–648. 10.1093/HMG/DDZ318.

108. Yue, L., Jin, M., Wang, X., Wang, J., Chen, L., Jia, R., Yang, Z., Yang, F., Li, J., Chen, C., et al. (2022). Compound Heterozygous Variants in a Surviving Patient With Alkuraya-Kučinskas Syndrome: A New Case Report and a Review of the Literature. Front Pediatr 10. 10.3389/FPED.2022.806752/FULL.

109. Filatova, A., Freire, V., Lozier, E., Konovalov, F., Bessonova, L., Iudina, E., Gnetetskaya, V., Kanivets, I., Korostelev, S., and Skoblov, M. (2019). Novel KIAA1109 variants affecting splicing in a Russian family with ALKURAYA-KUČINSKAS syndrome. Preprint at Blackwell Publishing Ltd, 10.1111/cge.13472 10.1111/cge.13472.

110. Dobritsa, A., Naters, W. van, Warr, C., Neuron, R.S.-, and 2003, undefined Integrating the molecular and cellular basis of odor coding in the Drosophila antenna. cell.comAA Dobritsa, WG van Naters, CG Warr, RA Steinbrecht, JR CarlsonNeuron, 2003•cell.com.

111. Lee, T., and Luo, L. (2001). Mosaic analysis with a repressible cell marker (MARCM) for Drosophila neural development. Preprint at Elsevier Ltd, 10.1016/S0166-2236(00)01791-410.1016/S0166-2236(00)01791-4.

112. Chen, J., Stork, T., Kang, Y., Sheehan, A., Paton, C., Monk, K.R., and Freeman, M.R. (2022). The Tre1/S1pr1 phospholipid-binding G protein-coupled receptor signaling pathway is required for astrocyte morphogenesis. bioRxiv, 2022.09.15.508188. 10.1101/2022.09.15.508188.

113. Sethi, S., and Wang, J.W. (2017). A versatile genetic tool for post-translational control of gene expression in Drosophila melanogaster. Elife 6. 10.7554/ELIFE.30327.

114. Li-Kroeger, D., Kanca, O., Lee, P.T., Cowan, S., Lee, M.T., Jaiswal, M., Salazar, J.L., He, Y., Zuo, Z., and Bellen, H.J. (2018). An expanded toolkit for gene tagging based on MiMIC and scarless CRISPR tagging in Drosophila. Elife 7. 10.7554/eLife.38709.

115. Li, T., Fu, T.M., Wong, K.K.L., Li, H., Xie, Q., Luginbuhl, D.J., Wagner, M.J., Betzig, E., and Luo, L. (2021). Cellular bases of olfactory circuit assembly revealed by systematic time-lapse imaging. Cell 184, 5107–5121.e14. 10.1016/J.CELL.2021.08.030.

116. Li, T., and Luo, L. (2021). An Explant System for Time-Lapse Imaging Studies of Olfactory Circuit Assembly in Drosophila. J Vis Exp 2021. 10.3791/62983.

117. Mirdita, M., Schütze, K., Moriwaki, Y., Heo, L., Ovchinnikov, S., and Steinegger, M. (2022). ColabFold: making protein folding accessible to all. Nature Methods 2022 19:6 19, 679–682. 10.1038/s41592-022-01488-1.

118. Emsley, P., and Cowtan, K. (2004). Coot: model-building tools for molecular graphics. Acta Crystallogr D Biol Crystallogr 60, 2126–2132. 10.1107/S0907444904019158.

119. Pettersen, E.F., Goddard, T.D., Huang, C.C., Meng, E.C., Couch, G.S., Croll, T.I., Morris, J.H., and Ferrin, T.E. (2021). UCSF ChimeraX: Structure visualization for researchers, educators, and developers. Protein Sci 30, 70–82. 10.1002/PRO.3943.

120. Goddard, T.D., Huang, C.C., Meng, E.C., Pettersen, E.F., Couch, G.S., Morris, J.H., and Ferrin, T.E. (2018). UCSF ChimeraX: Meeting modern challenges in visualization and analysis. Protein Sci 27, 14–25. 10.1002/PRO.3235.

121. Fraser, A.G., Kamath, R.S., Zipperlen, P., Martinez-Campos, M., Sohrmann, M., and Ahringer, J. (2000). Functional genomic analysis of C. elegans chromosome I by systematic RNA interference. Nature 408, 325–330. 10.1038/35042517.

122. Livak, K.J., and Schmittgen, T.D. (2001). Analysis of relative gene expression data using real-time quantitative PCR and the 2(-Delta Delta C(T)) Method. Methods 25, 402–408. 10.1006/METH.2001.1262.

